# Mapping the Multiscale Organisation of Escherichia Coli Chromosome in a Hi-C-integrated Model

**DOI:** 10.1101/2020.06.29.178194

**Authors:** Abdul Wasim, Ankit Gupta, Jagannath Mondal

**Affiliations:** Tata Institute of Fundamental Research, Centre for Interdisciplinary Sciences, Hyderabad 500046, India

## Abstract

The chromosome of Escherichia Coli (E. coli) is riddled with multi-faceted complexity and its nature of organization is slowly getting recognised. The emergence of chromosome conformation capture techniques and super-resolution microscopy are providing newer ways to explore chromosome organization, and dynamics and its effect on gene expression. Here we combine a beads-on-a-spring polymer-based framework with recently reported high-resolution Hi-C data of E. coli chromosome to develop a comprehensive model of E. coli chromosome at 5 kilo base-pair resolution. The model captures a self-organised chromosome composed of linearly organised genetic loci, and segregated macrodomains within a ring-like helicoid architecture, with no net chirality. Additionally, a genome-wide map identifies multiple chromosomal interaction domains (CIDs) and corroborates well with a transcription-centric model of the E. coli chromosome. The investigation further demonstrates that while only a small fraction of the Hi-C contacts is dictating the underlying chromosomal organization, a random-walk polymer chain devoid of Hi-C encoded contact information would fail to map the key genomic interactions unique to E. coli. Collectively, the present work, integrated with Hi-C interaction, elucidates the organization of bacterial chromosome at multiple scales, ranging from identifying a helical, macro-domain-segregated morphology at coarse-grained scale to a manifestation of CIDs at a fine-grained scale.

## INTRODUCTION

The folding of the 4.64 Mbp circular chromosome, with a contour length of 1.6 mm, inside 2-4 *µ*m long spherocylindrical cell(1, 2) of *Escherichia coli* (*E. coli*) (∼1.5*µm*^3^ in volume) is a complex process, mediated by numerous factors and cues. Several decades’ investigations on chromosomal DNA of *E. coli* have rendered a picture of a highly condensed form called the nucleoid which is a dynamic macromolecular complex of the genetic material and nucleoid associated proteins (NAP) along with proteins such as RNA Polymerases(RNAPs)(3). The multitude of experiments on *E. coli* chromosome have highlighted a ring-like architecture of nucleoid whose organization results from a combination of processes including DNA supercoiling(4), NAP-induced condensation of DNA(5), crowding and non-equilibrium processes like transcription(3). In rapidly growing *E. coli* cells, nucleoid is severely compacted at the centre of cytoplasm with ribosomes being strongly concentrated at the periphery of the nucleoid(6). The multiscale organization underlying the *E. coli* nucleoid is only slowly getting recognised.

A set of classic experiments, including fluorescence in situ hybridisation (FISH) techniques (7, 8) and site-specific recombination assays(9) had indicated spatial proximity and increased interactions among genetically distant DNA sites, giving birth to the idea of organization into a ring-like chromosomal architecture comprised of four large macrodomains, namely Ori, Ter, Left, Right and two Non-Structured regions-Right (NS-R) and Left (NS-L). These macrodomains are considered to be spatially segregated from each other(7, 8). Another investigation via fluorescence labelling of the genetic loci had proposed a model in which *E. coli* nucleoid displays a linear order of the locus distribution along the axial dimension of spherocylinder(10). More intriguingly, recent experimental studies brought into light the helical folding of the chromosome(11, 12). In particular, it was found that the circular chromosome twists along the long axis to form a helix like, achiral conformation. The complexity underlying bacterial chromosome has motivated a series of computer simulation studies and models to describe the architecture of *E. coli* chromosome. Some of them are based on coarse-grained models(6, 13, 14, 15, 16), while others used a more fine-grained approach for bacteria with base pair or approaching base-pair resolutions(17, 18, 19, 20). While these models significantly contribute to our current understanding on *E. coli* chromosome, most of these models are phenomenological in nature and lack attempts in integrating experimental data.

In the first data driven model for *E. coli* chromosome, ChIP-chip data for RNAP was integrated into a fine grained polymer model(21). On the other hand, the emergence of high resolution chromosome conformation capture data in bacterial cells (22, 23, 24) has helped detect the interaction frequency between any two genomic loci in the whole genome of an organism and build data-informed models of bacterial chromosomes. Hi-C has the unique ability to determine the interaction frequency map of whole genome of an organism at high resolution(25). The ensemble nature of the resulting contact frequency matrices automatically incorporate the inherent stochasticity of the chromosome’s interactions. Using Hi-C and super-resolution microscopic imaging the 3D chromosome conformation of *Mycoplasma pneumoniae* was determined(24). Similarly a Hi-C driven model was proposed for investigation of conformational dynamics of *Caulobacter cresentus* chromosome(26). All of the above data driven approaches show that with emerging experimental techniques, one can enrich a computational model via integrating it with experimental data. Such theoretical studies can introduce specificity such as macrodomain formation, their locations and relative sizes for a bacterium into its model(s).

In this regard, the recent report of high resolution Hi-C interaction maps of *E. coli* chromosome, has been a key breakthrough(22). It brings out the salient features of NAP-mediated multiscale organization underlying the *E. coli* chromosome architecture. It also opens up promising opportunities for developing high resolution and quantitative model for the same. In the current work, we present computer simulations of the *E. coli* chromosome by integrating beads-on-a-spring polymer model with recently reported Hi-C interaction matrix of *E. coli* chromosome(22). As would be detailed in the article, the model, developed at a 5 kbp resolution, unifies multiple existing hypothesis related to *E. coli* chromosome’s architecture and demonstrates long-range organization into multiple macrodomains. In addition, the presented Hi-C integrated chromosome model unifies a wide array of experimental data such as FISH data(7), recombination assay(9) and ChIP-chip data based simulations. As would be revealed in the text that follows, the model vividly manifests the multiscale and multi-faceted organization of bacterial chromosome, namely a helical, macrodomain separated morphology at coarse-grained scale and CIDs at a fine-grained scale. (21)

## MATERIALS AND METHODS

Figure 1 outlines the schematic of the integrative method used in the current work to generate the chromosome structures via combination of beads-on-a-spring model and Hi-C contact matrices. Below we detail different segments of the methods employed in the current work.

**Figure 1.**
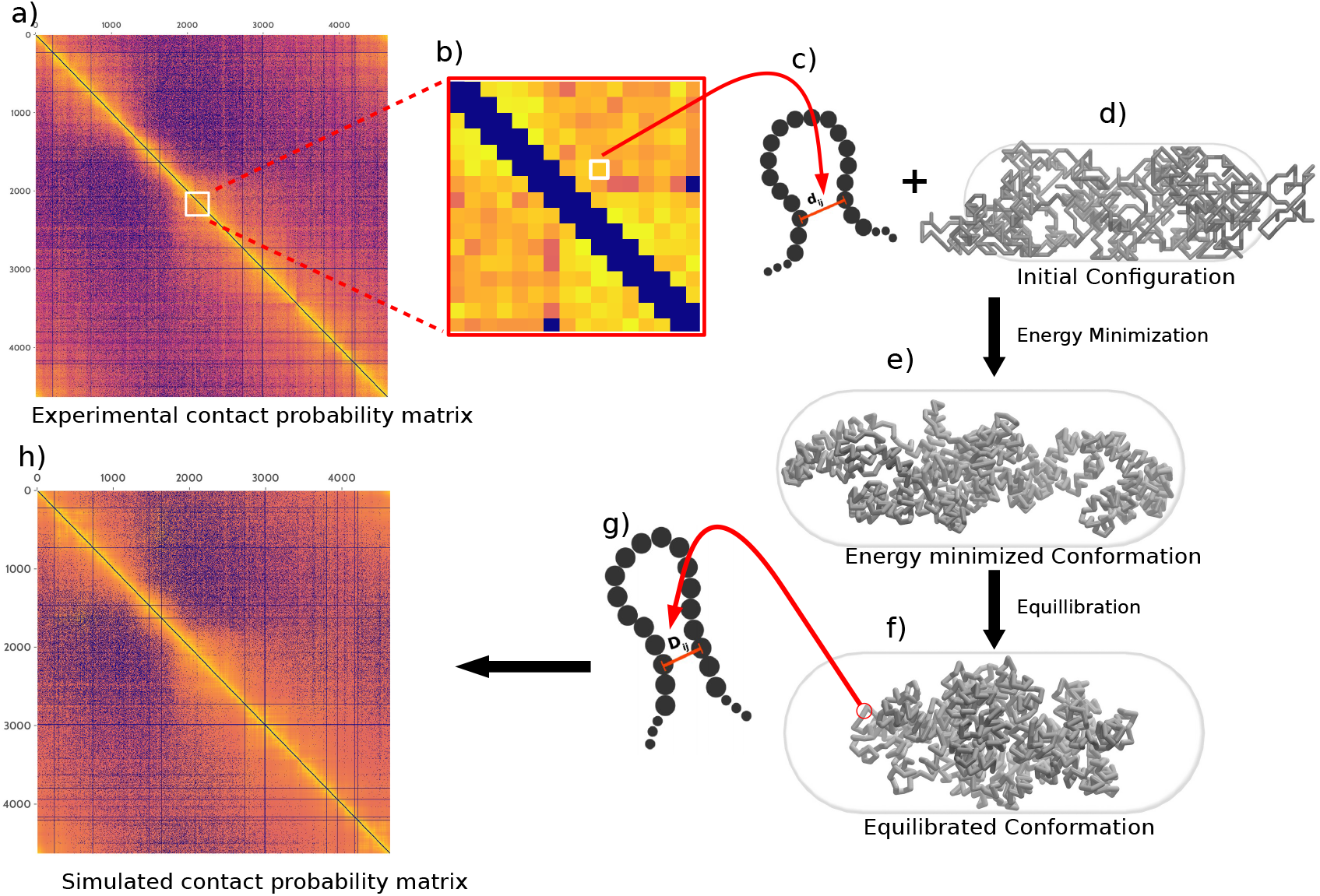
A schematic of the method used to generate an ensemble of final chromosome structures. **a)** Contact probability matrix from Hi-C experiment. **b)** Expanded section of the matrix. **c)** The experimental contact probabilities are mapped onto bond distances and bond strengths. d_*ij*_ is the initial bond between particles i and j. **d)** The calculated distances were implemented on a beads-on-a-spring model (having only repulsive Lennard jones potential initially) as harmonic constraints. **(e)** An energy minimized conformation with interactions as described previously. **f)** Chromosome after equilibration of an energy minimized conformation. **g)** D_*ij*_ is the equilibrium bond distance between i and j. Average bond distances from equilibrium conformations were converted to a contact probability matrix. **h)** Hi-C contact probability matrix from simulation. The simulated and the experimental matrices are compared and the simulated matrix is used for further analysis.

### Hi-C data processing

We used the data made available by *Lioy et al. (2018)* (22) with the GEO accession number *GSE*107301. SRA (Sequence Read Archive) files were splitted into both reads of pair-end sequences using *fastq-dump*. We used hiclib python library (https://bitbucket.org/mirnylab/hiclib) provided by Leonid Mirny’s group for further processing of the fastqs. Using hiclib, iterative mapping was performed with a minimum sequence length of 20bps and a step length of 5bps. Alignment reads were saved in BAM (Binary Alignment Map) files and processed according to 5kbp resolution and filtered using hiclib default fragment level filtering functions, namely: duplicates, large, extreme, dangling ends, etc and they were binned into 928 bins according to 5 kbp resolution. Further bin level filtering was performed to remove low coverage bins, bins with only small area sequenced and diagonal and adjacent to diagonals bins. Then the raw matrix was extracted using h5py python library from HDF5 file generated by hiclib library, we also set the two extreme elements on the off-diagonal of the matrix to zero, since they also are reads from adjacent regions of the chromosome, due to the chromosome being circular. The matrix is then normalised by using sequential component normalization (SCN) (27) in which, first all the column vectors were normalized to one using euclidian norm followed by each row vector and the whole process was iterated until the matrix become symmetric again (3 iterations in our case). The normalized matrix is converted to a contact probability matrix by dividing each row by its maximum value (28) followed by a resymmetrization of the matrix(Figure 1a).

### Model details and interaction potentials

We modelled the 4.64 × 10^6^ bp *E. coli* chromosome as a beads-on-spring polymer with each bead representing a 5 × 10^3^ bp (i.e. 5 kbp) nucleotides, which is the resolution of Hi-C interaction maps. The 5 kbp nucleotides are indexed and annotated as per the genetic sequence of wild type *E. coli* MG1655 (GenBank ID: U00096.2) and modelled as non-overlapping van der Waals particles, giving rise to a total number of 928(= 4.64 × 10^6^*/*5 × 10^3^) beads. The polymer beads are subjected to a spherocylindrical confinement commensurate with average dimension of *E. coli* at 37°C in LB media (axial length (including end-caps) of 2.482 *µ*m and the diameter of 0.933 *µ*m (1)) (See figure S1). As detailed in the *SI Methods*, based on an approximate volume fraction of the chromosome of 0.1(29) (with respect to cell volume) and the spherocylindrical confinement dimensions, the individual bead diameter (*σ*) was determined to be 0.067 *µ*m. The non-bonded interactions are purely repulsive and are given by the repulsive part of Lenard-Jones potential 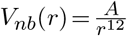 where, A = 4*ϵσ*^12^. For the simulations, A=1.0 kJ mol^*−*1^ *σ*^12^ has been used.

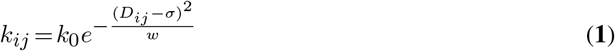

Adjacent beads of the polymer are connected by strong (300 kJ mol^*−*1^*σ*^2^) harmonic springs with *σ* as the equilibrium bond length. Hi-C contacts are also modelled as harmonic springs but with distance-dependent force constants and probability-dependent bond lengths(Figure 1b). For Hi-C contact probability matrix *P*, we define the distance matrix, *D* as 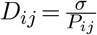, where *ij* suggests the element in the i^th^ row and j^th^ column of the matrices. The restraining potential between a pair of Hi-C contacts at a separation of *r*_*ij*_ is given by 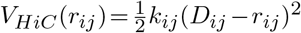. It should be noted that the matrix *P* is a sparse matrix with large number of the probability matrix elements being close to zero. Therefore a lot of the elements in D would translate to large distance close to ∞. To take this into account, in our simulations, the force constants for the bonds incorporating the Hi-C contacts have been modelled as a gaussian function of the Hi-C distances (Eq.(**1**)) (as represented in Figure 1b and 1c). It should be noted that the choice of using Eq. (**1**) is not arbritrary. We have also explored other possible equation (Eq. (S5)) in place of Eq. (**1**) (see *SI Interaction potentials* section), but we found Eq. (**1**) to be a better function for modelling (see *SI text* and figure S2-S5 for details).

Here *k*_0_ is an amplitude term that determines the upper limit to the force constants of the “Hi-C bonds”. This function essentially implies weaker values of force constant for larger distances. This function naturally takes *V*_*HiC*_ (*D*_*ij*_)= 0 for *D*_*ij*_ = ∞. In equation S7 in *SI methods, k*_0_ and *w* are parameters that need to be optimised. The metric used to optimize *w* is a Pearson correlation coefficient between the experimental and the *filtered* simulated contact probability matrices (filtering has been performed as explained in *SI Materials and Methods*).

To implement the effect of spherocylindrical confinement induced by a *E. coli* cell, a restraining potential (Eq. 2) has been used

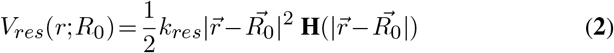

**H** is a step function and gets activated only if any chromsome bead attempts to get out of the spherocylindrical confinement. *R*_0_ is the center of the spherocylinder. *k*_*res*_ determines extent of elasticity of the cell boundary. For simulations we have used 310 kJ mol^*−*1^*σ*^2^.

### Simulation details

All simulations are performed using open source package GROMACS 5.0.7(30). The source code of the program was modified by us to implement the interaction potential function of the spherocylindrical confinement. All other bonded and non-bonded interaction potentials were introduced by using default GROMACS utilities. At the start of all simulations, a 928 bead polymer is generated bead-wise (using home made python scripts), avoiding any inter-bead overlap. The generated conformations are subsequently energy-minimized by steepest descent algorithm using GROMACS(Figure 1d and 1e). The energy minimized polymer is then subjected to Langevin dynamics in NVT ensemble (Figure 1e and 1f). The temperature of the system was maintained using Langevin thermostat at 303<u>K. T</u>he time step for equilibration or production run is 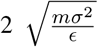, where m is the mass of 1 bead (see GROMACS manual for units). Each production run has been run for 2 × 10^6^ steps from which only the last 2000 frames have been used for further analysis. 200 simulations have been performed with independently generated initial structures to capture the stochastic effect of the ensemble. Using the last 2000 frames of each individual simulation, we generate a distance matrix and a probability matrix, whose values have been averaged over the number of frames of that particular simulation. Subsequently, we generate the final simulated distance matrix and the probability matrix by averaging over frame averaged matrices over all 200 independent simulation (represented by Figure 1f, 1g, and 1h).

We had also performed simulations with 12 × 10^6^ steps to check for proper equilibration of the polymer. On comparing the simulated contact probability matrices for simulations with 2 × 10^6^ and 12 × 10^6^ steps (Figure S7), we concluded that the polymer has equilibrated within 2 × 10^6^ steps and this simulation length will be sufficient for investigation.

## RESULTS AND DISCUSSION

### Simulations reconstruct experimental Hi-C data

Figures 2a and 2b compare the experimental Hi-C contact probability matrix of *E. coli* chromosome with that obtained from our simulations. A single intense diagonal indicates a smaller distance with a higher contact probability between neighbouring chromosomal regions. Absence of a secondary diagonal in the Hi-C interaction matrix reflects the lack of contacts between the two replication arms of the chromosome. A characteristic feature of prokaryotic chromosome is its circularity. High contact probabilities at the end regions of the Hi-C interaction matrix in Figure 2b (right most upper and left most lower corner) assure circularity of the chromosome in the current model as well.

**Figure 2.**
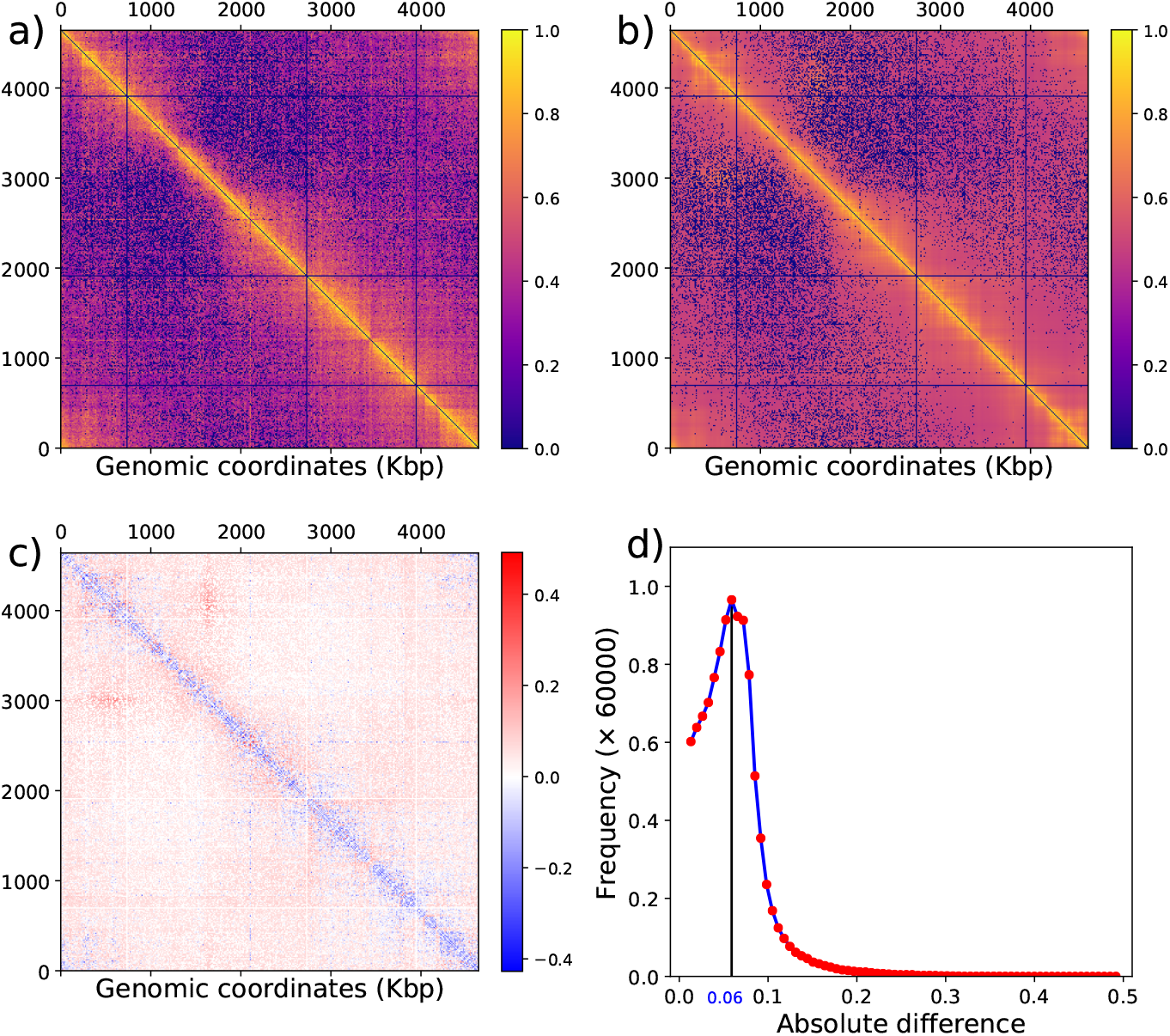
**a)** Heatmap of experimental and **b)** simulated (filtered) Hi-C matrix. **c)** Heatmap of difference between contact probability matrix (simulated - experimental matrix), negative number or blue color indicates higher probability value in experimental matrix and positive number or red color indicates higher probability in simulated matrix. **d)** Distribution of absolute differences between the two contact probability matrices(experimental and simulated).

A Pearson correlation coefficient of 0.89 between experimental and simulated probability matrix indicates a very good agreement between simulated and in-vivo chromosome conformations. For a direct comparison, we also plotted the heatmap of the difference between the experimental and simulation-derived contact probability matrices (Figure 2c) and the resulting histogram of absolute values from the heatmap (Figure 2d). The difference heatmap shows that for smaller genomic distances (i.e. bins near the diagonal) the contact probability is relatively higher in experimental matrix (blue regions). For rest of the matrix the difference is negligible. The distribution of absolute values from the heatmap in figure 2c also shows that the disagreement between experiment and simulation is extremely small as the major difference is less than 0.1. Together these analysis suggest a good correspondence between experimental and simulated Hi-C matrices. Thus our model is robust for three-dimensional reconstruction of the *E. coli* bacterial chromosome and can be explored for investigating and predicting key features of the chromosome at multiple length-scale.

### Illustration of Macrodomain organization in the simulated chromosome conformation

Figure 3a is a representative configuration of the *E. coli* chromosome, as obtained from current simulations. Multiple experimental investigations involving *E. coli* chromosome have, in the past, proposed the existence of a set of macrodomains in its genome (7, 8, 9). Accordingly, in our model, we have color-coded the beads as per the annotation of proposed genetic sequences of chromosome macrodomains. The snapshot in Figure 3a shows that all the annotated macrodomains are well segregated from each other. The segregation of macrodomains along the long axis can also be inferred from the average densities of 4 macrodomains and 2 non-structured regions, shown in Figure 3c.

**Figure 3.**
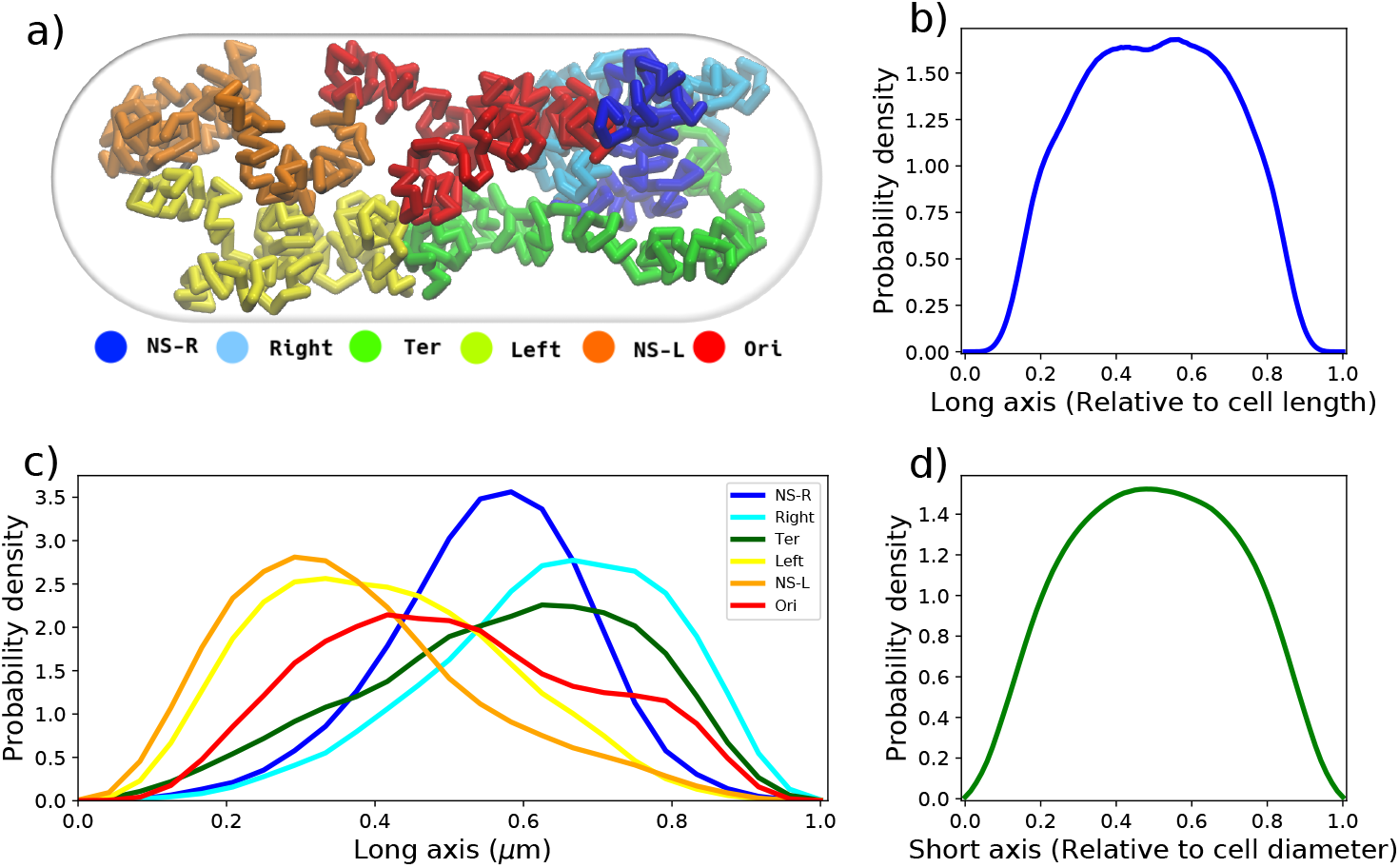
**a)** Snapshot of the equilibrated chromosome from a trajectory with macrodomains colored and cell dimensions as mentioned in methods. **b)** Linear density of chromosome along long axis. **c)** Average density of each macrodomain and non-structured regions in 200 trajectories and 2000 frames for wildtype *E. coli* with respect to the cell’s long axis. **d)** Linear density of chromosome along the short axis.

The Left and Right macrodomains have occupied cell’s left and right halves which was also found in earlier experiments(31). Specifically, Ori (red) and Ter (green) macrodomains are aligned at the mid-cell, as visible from the structure in Figure 3a and from their respective linear densities (Figure 3c). Although the position of all the macrodomains are variable across replica of trajectories, the densities in Figure 3c suggest that there is effectively an average positional preference for each of the macrodomains inside the cell: Mid-cell for Ori and Ter, left and right half of the cell for Left and Right macrodomains respectively. Interestingly, the representative snapshot also suggests that the chromosome is not completely spread across the entire axial dimension.

We also calculated the average linear density profile for the whole chromosome along both long and and short axes (z- and x-respectively), shown in Figure 3b and d. The chromosome occupies the central region along the long axis without any external restrictions as found in previous experimental and theoretical studies(6, 32).

### Capturing a linearly-organised genomic architecture and positioning of oriC and diff of nucleoid

Precedence of FISH data for the inter-focal distances between various loci throughout *E. coli* chromosome(7) allowed us to compare distances from our simulated model with FISH-based measurement. A Pearson correlation coefficient of 0.83 (Figure 4a) shows reasonably good agreement between distance calculated from our model and FISH-determined inter-focal distances. Since the distances are calculated in real units (*µm*), the slope of the fit gives us the relative size of the simulated chromosome with the in-vivo counterpart. In Figure 4a, a slope of 1.35 indicates that the loci distances are larger in the current model than that found in experimental setup.

**Figure 4.**
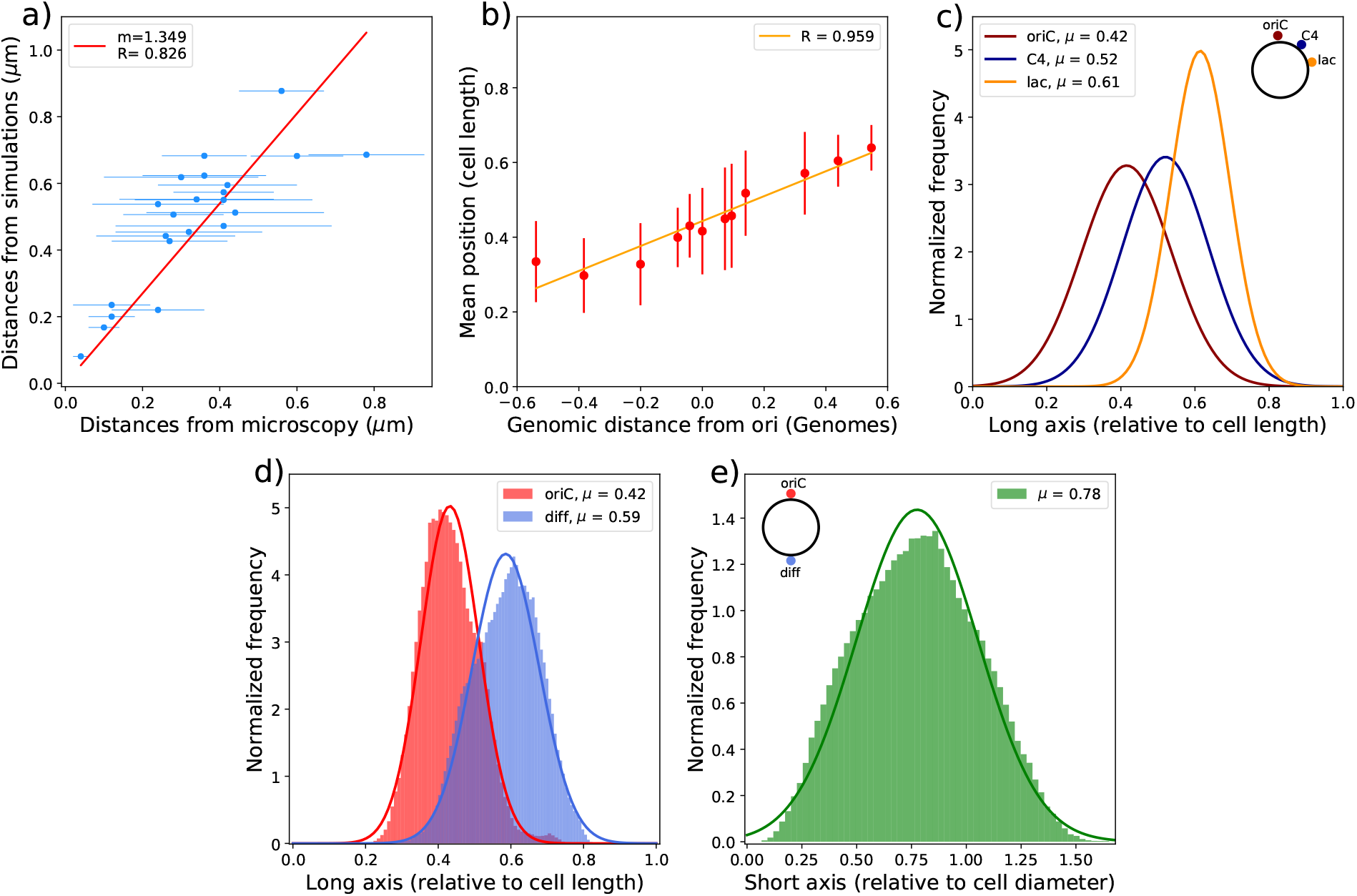
**a)** Distances measured from our simulations (y-axis) for Hi-C data of *E. coli* at 37 °C in LB media vs. distances measured via FISH (x-axis), measured at 25 °C in Minimal media. **b)** Mean position along long axis vs. genomic distance from oriC as mentioned by *Wiggins et al. (2010)*(10). **c)** Distribution of position of three loci: oriC, C4 and lac along the long axis(Z-axis). **d)** The distribution of oriC and diff locus with respect to cell length where 0.5 implies the midcell position. **e)** The distribution of distance between oriC and diff locus in 200 simulations and 2000 frames each.

We speculate that this might have to do with the difference in media and temperature between FISH (25°C in minimal media) and Hi-C (37°C in LB media) measurements.

For a more comprehensive investigation, we determined the position-distribution of a few loci (genomic locations obtained from *Wiggins et al*.(10)) along the long axis of cell and plotted their mean position (averaged over all trajectories) as a function of their genomic location (relative to that of oriC). As depicted by the linear fit of Figure 4b, we find that the linear organization of most of the loci along long axis of cell is also retained in the present model of chromosome. However, the Hi-C based model suggests that the separation between the loci is smaller than that reported in the experimental investigations(10) because the fitted line is steeper in the experimental data(10) than our current model. These observations imply that though such elaborate loci arrangement is more evident in slow growing cells, they are also retained in rapidly dividing cells (the present scenario), albeit to a smaller extent.

In a previous fluorescence based assay(10), positions of multiple genetic loci in *E. coli* cells were monitored. From their spatial positioning a ‘linearly organised’ architecture of the chromosome was proposed. In particular, this investigation hypothesised that these loci are linearly positioned along the long axis of the chromosome. To test this hypothesis in our Hi-C encoded polymer model of chromosome, we plot the distribution of three loci oriC, C4 and lac along long axis of cell(Figure 4c). (These are the same loci position measured in Figure 2A by *Wiggins et al. (2010)*). Our model predicts that these loci are linearly positioned along the long axis of the cell, in accordance with the experiment(10).

Two loci, namely, oriC and diff are known to play a pivotal role in *E. coli* chromosome division and segregation (8, 10, 33). OriC is the origin of replication from where replication of chromosome starts and diff is another locus present in the Ter macrodomain and opposite to oriC in genomic distance. Diff is the last region on the chromosome to be replicated after which the two chromosomes segregate. In exponentially dividing cells, these two loci have been hypothesized to be present at the mid-cell for majority of the time(31). To investigate the location of these two loci in our model, we calculated their average positional distribution. As we can see from Figure 4d, the mean position for oriC and diff are 0.42 and 0.59 in cell length along the long axis. The distribution of distance between oriC and diff (Figure 4e) shows that the average distance between the two loci is also approximately equivalent to the diameter of the cell. Together, these distributions show that these two loci are axially present at the mid-cell and radially apart by the cell’s periphery, which is an important feature of the rapidly dividing cells in case of *E. coli*.

### Macrodomain sizes and prediction of recombination assay data

To determine the approximate size of the macrodomains we calculated the radius of gyration (*R*_*g*_) for each macro-domain (Figure 5a). Interestingly, we find that *R*_*g*_ s of these domains are matching with *R*_*g*_ s predicted by *Hacker et al*.’s oriC@midcell plectonemic model (21), which is in accordance with our earlier prediction of mid-cell location of oriC loci. We find that Ter has the highest R_*g*_ (0.35*µ*m) among all macrodomains and Non-Structured regions (see Table S1). This is because in most of the structures, unlike other macrodomains, Ter was present as a linear chain, as evident by its highest end-to-end distance (Table S1). This result is also consistent with the model proposed by *Wiggins et al.(2010)*(10) based on fluorescence-based experiments, that the Ter region acts like a linking part between Right and Left macrodomains which are spanning the right and left cell halves respectively.

**Figure 5.**
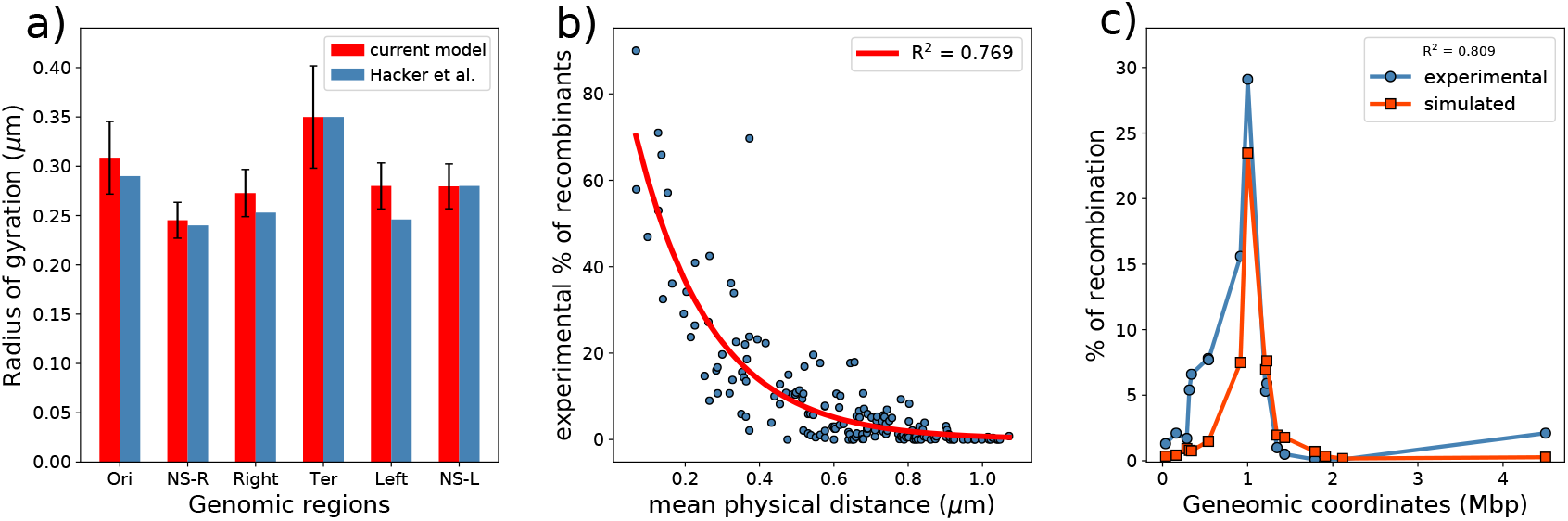
**a**) Average radius of gyration of each macro-domain from current model as compared to that given by *Hacker et al. (2017)*(21) for oriC@midcell plectnomic model. **b)** Comparison between the recombination frequencies provided by *Valens et al. 2004* (9) vs. mean physical distance between recombination loci and red line indicates the single exponential fit. **c)** A representative plot of recombination frequencies predicted by mean physical distance, blue: experimental data and red: predicted by simulated data.

The model is able to predict the recombination assay percentages. We first obtained an exponential fit between experimental percentage of recombinants and distances calculated for such loci pairs from simulations (Figure 5b). Using the fit, we calculated the percentage of recombination for reported pairs of loci(9) and plotted the percentages with respect to their genomic distances (Figure 5c), similar to a previous investigation(21). The fit suggests a good correlation with *R*^2^ value of 0.769. Figure S8 shows the predicted recombination for other six loci experimented (9) which are also in good correlation with the experimental data.

### Helicoid conformation of the chromosome

Recent experimental investigations(11, 26) have hypothesised that the bacterial chromosome, most likely, adopts a helical conformation. To explore any likely features of the helical conformation of the chromosome from our model, we plotted the centre of geometry(COG) of a set of beads (Figure 6a and Figure S9) ranging from *i*-10 to *i*+10, where *i*=1 to 928 and calculated its crossing number and writhe. Crossing number is defined as the number of times the chromosome intersects itself when the 3D conformation is projected onto a 2D plane. When such crossings are viewed in 3D, they actually do not intersect. In 3D, if one looks closely at two segments that intersect each other on a plane, then it would be seen that one segment goes over or under the other segment. Depending on whether it goes over or under, that crossing is given a sign, + or - respectively. The sum of all crossings with its sign is called the writhe (see *SI Methods* for detailed discussion on writhe). Upon calculation of the average number of crossings for the COG contours of the last 2000 frames, we found an average crossing number of 1.522 with an average writhe of 0.046. Together, these numbers signify that though the COG contour of the chromosome crosses itself a couple of times, the probabilities of over- and under-crossings are equal. This makes the COG contour resemble a twisted circular ring, as seen from Figure 6a.

**Figure 6.**
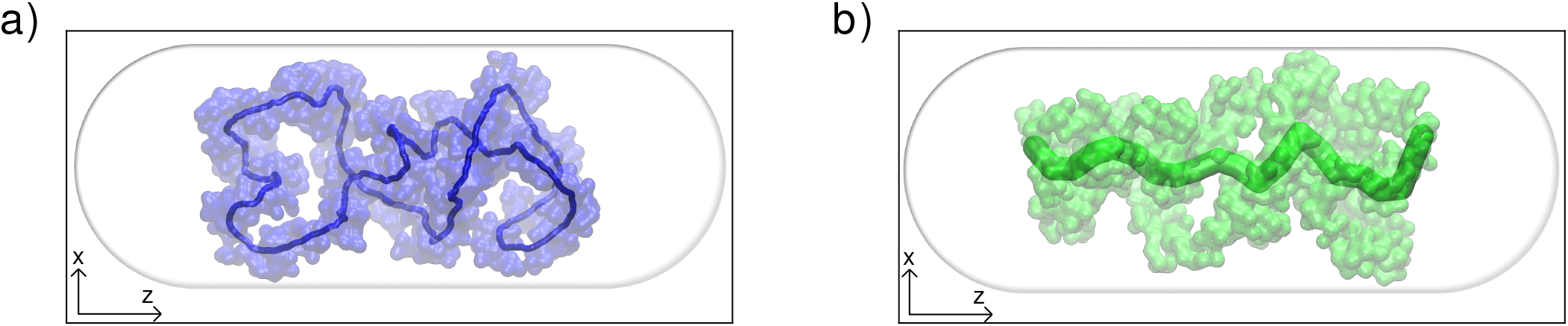
**a)** A representative image of the contour of the polymer generated by plotting the centre of geometries of a set of beads (21 for the image provided here). **b)** Smoothed centre of geometries for the chromosome viewed from the xz plane.

To corroborate predictions from our model with experimental observations by Marko and coworker (11), we divide the cell along the long axis into 20 equal slices (referred to, from now on, as z-slicing) and plotted the COGs of each slice (Figure 6b). We find that the COGs obtained from the z-slices in our model are similar to those experimentally calculated(11). By fitting the coordinates obtained from z-slicing to a polygon, we calculated it’s average writhe, as described by Klenin *et. al*(34). We find that the writhe is equal to -0.092 ±3.066. The difference in growth conditions for the bacteria in the current study and the experimental study(11) might result in the slight mismatch between the values. But we also get a very low mean writhe which suggests that in our model too the chromosome tends to form a helicoid with no bias to arrange itself in a clockwise or anti-clockwise fashion, thus no net chirality.

We also calculate the average number of crossings and the average writhe for the whole chromosome using the raw coordinates we obtain from our model. The crossing number and writhe for the whole polymer was found to be 923 and 0.49 respectively (Figure S10). These values of writhe and crossing numbers emphasizes that the circular chromosome must have adopted a twisted ring like conformation which we already saw in Figure 6a. Values of the writhe close to zero signifies that the chromosome, which adopts a helix like conformation, has equal probabilities of over- and under-crossings, which is also a characteristic feature of a helical structure.

### Insights on global packing of the Chromosome

A common approach for exploring global packing of chromosome is the investigation of scaling of contact probability with respect to genomic distance(25). Earlier, based upon similar analysis of intra-chromosomal contact probability as a function of genomic distance, a power law scaling of contact probability between 500Kb and 7Mb for human chromosome was predicted(25). The contact probability was found to be scaled as the inverse of genomic distance (∼ *s*^*−*1^), which is in agreement with a previously proposed attribute that chromosomes are fractal in nature(35).

Fractal polymers are knot free which is important for the segregation of daughter chromosomes during cell replication(25, 36). For *E. coli*, similar analysis of the intra-genome contact probability as a function of genomic distance, as extracted from experimentally determined Hi-C matrix, produces a scaling factor of ∼−0.77, suggestive of deviation from a perfect fractal. In our current model (Figure S11a) the contact probability scales as ∼*s*^*−*0.55^ in 10Kb to 1Mb range. On the other hand, scaling of root-mean-squared (RMS) end-to-end distance with genomic distance suggests otherwise (Figure S11b). RMS end-to-end distance scales as *s*^0.36^ which is close to the expected value for scaling of RMS end-to-end distance with genomic distance for a fractal globule polymer which is *s*^1*/*3^.

To confirm the fractal nature we explored the presence of knots in our ensemble of structures explicitly using an external python library(37). Knot analysis revealed that all conformations, whose initial structures did not have any knots, are knot free. Overall, 86% of the total conformations, including the ones which started out with knots, were knot free. This implies that the chromosome is not an equilibrated globule, although it may not fold in a completely fractal way.

### Insights on local packing of the Chromosome

Figure 7a shows the CID boundaries calculated from the experimental contact probability matrix. The boundaries have been calculated using the Directionality Index (DI) algorithm with a window size of 100 Kbp (Figure S12). We see from Figure 7b that DI on the simulated contact probability matrix provides us with a lesser number of peaks. Therefore to calculate the boundaries directly from the structure, we developed a method which we call the R_*g*_ map method (see *SI Methods*). In this method we use a moving window averaging of radius of gyration of continuous segments of chromosome. This approach gives rise to a ‘radius of gyration map’ (R_*g*_ map). Using this approach, we calculated the radius of gyration of fine-grained segments for each bead (for example: for *n*_*th*_ bead: the R_*g*_ will be calculated for 20 beads starting from *n* − 10^*th*^ to *n*+9^*th*^ bead) for all the 928 beads in the chromosome. We have calculated the R_*g*_ map at four different window sizes: 10, 20, 50, 100 beads, corresponding to 50, 100, 250, and 500 kbp genomic segments respectively, to investigate the features of our model pertaining to the local density of chromosome at different scales. This method also enables us to probe the variation in local density of the chromosome along it’s contour.

**Figure 7.**
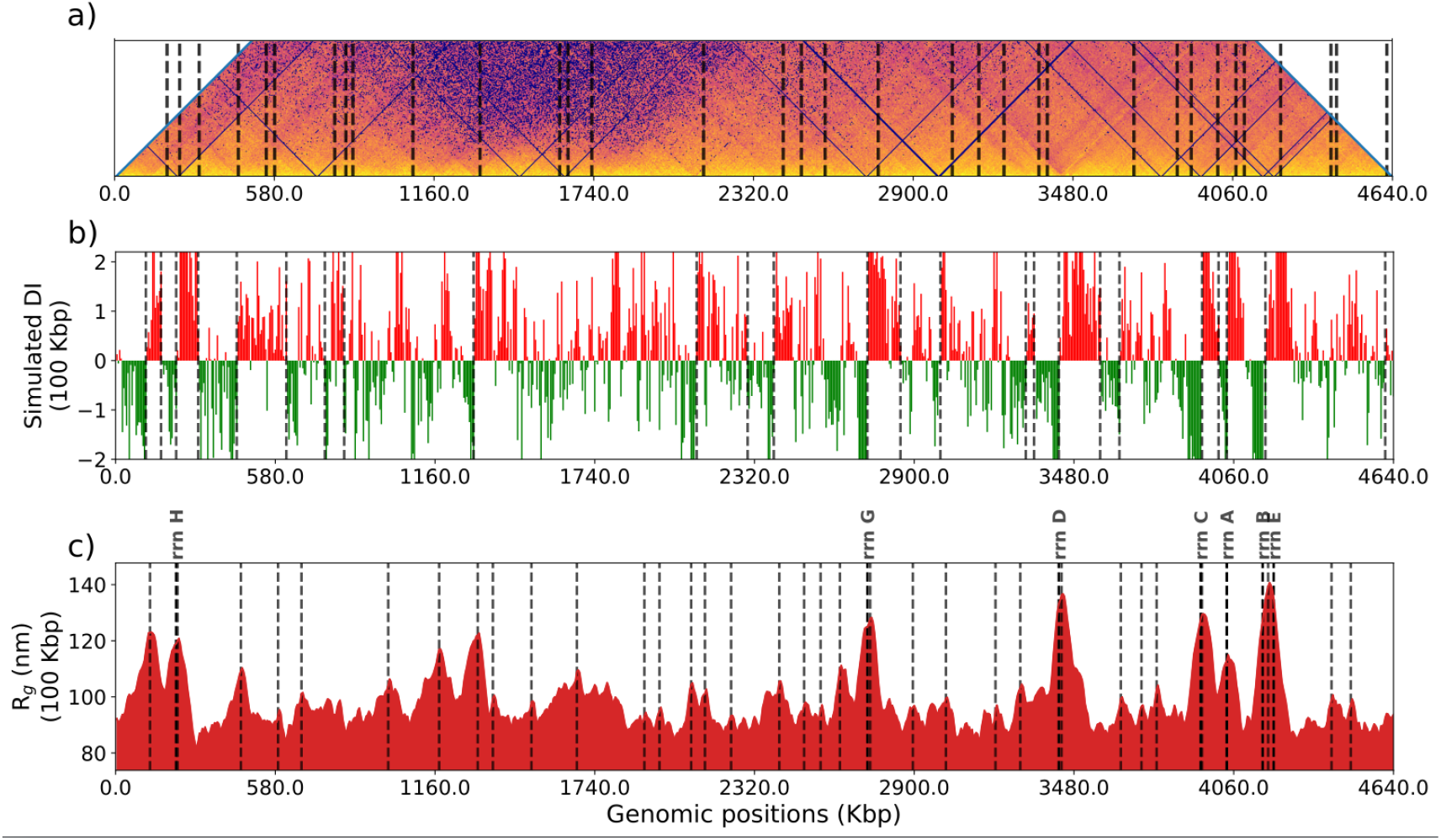
**a)** The main diagonal of the average contact probability matrix rotated 45° clockwise with black vertical lines showing the position of CID boundaries. **b)** Directionality index (100kb) calculated from simulated matrix for wildtype *E. coli* (MG1655 in LB at 37°C in exponential growth phase) black vertical lines shows the CID (chromosome interaction domains) boundaries, (calculated as given by *Le et al.(2013)*(23)). **c)**Radius of gyration map for wildtype *E. coli* (MG1655 in LB at 37°C in exponential growth phase) with respect to genomic coordinates with a moving window size of 20 beads (100kbp), black vertical lines represent the peaks in the map and rrn operon sites respectively

Figure 7c shows the Rg map for wild type *E. coli* (grown at 37°C in LB) at a window size of 20 beads or 100 kbp region. The black vertical lines, in the Rg map, highlight the most prominent peaks which were calculated with the help of a peak caller from scipy(38), which is tuned to get the exact number of peaks as predicted by directionality index (DI) at 100 kbp size. We report a correlation of 0.99 between the positions of the CID boundaries detected by R_*g*_ map and DI (Figure S13). Previously it was shown that the DI boundaries, when visualised with the heat map of matrix, appear at the vertices of the triangles along the diagonal of the matrix. Therefore, we also compared our Rg map with the diagonal of the heat map (shown in Figure 7a). The peak positions calculated from the Rg map are also marked in the Figure 7c and qualitatively we can see that these peaks are present at the vertices of triangles in most of the cases. Therefore, the comparison in Figure 7 suggests that our model is capable of capturing local chromosomal structures such as chromosomal interaction domains (CID). Together, Rg map comes out to be an effective tool in unravelling the local structure from our simulation model.

In a study on *Caulobacter crescentus*, it was found that, generally, one or many highly expressed genes were present at the CID boundaries(23). In rapidly dividing cells, the rrn operons are expressed at a higher rate than other metabolic genes due to the requirement of ribosomes for protein synthesis(39). Since the Hi-C experiment was also performed on rapidly dividing cells, high expression of rrn operons was expected. In another study, rrn operons showed higher transcriptional propensity, measured as RNA/DNA ratio, and overlap with CID boundaries(40). Therefore, motivated by previous experiments (23, 39, 40) and good agreement between peaks in R_*g*_ map and CID boundaries, we compared the peaks in R_*g*_ map with rrn operons’ genomic locations in *E. coli*. We found that six of the peaks in R_*g*_ map correspond to the genomic location of rrn operons as shown by bold, black vertical lines in Figure 7c.

Physically, higher R_*g*_ value indicates that the chromosome segment (in case of Figure 7c - 20 beads) is occupying more volume than the adjacent segments of same length. Similarly, lower R_*g*_ value implies that the segment is compact and occupies lower volume than its adjacent segments. Therefore our results imply that the local DNA density is lower in the vicinity of highly expressed genes, a phenomenon earlier observed in the eukaryotic cells(41). These results suggest similarity with the ‘transcription centric’ approach used in a previous model(21). These low density regions i.e. with high R_*g*_ value in R_*g*_ map are thus equivalent to plectoneme free regions(PFR)(21). Together, our model, via encoding Hi-C data, is able to capture all the spatial informations of the chromosome such as macrodomain structure, plectonemes, CIDs, and transcription details.

### Assessing the importance of the genomic contacts in chromosome organizations

While modelling the Hi-C contact probability data into distance restraints, we had made use of a sparse interaction matrix and considered only a fraction of high-contact probability values from the probability matrix (5.06% of the total number of contacts). To assess the importance of the small percentage of the Hi-C contact probability matrix that has been used as an input in the model, we carried out “control simulations” in which a self-avoiding polymer chain of same bead numbers were modelled but no Hi-C restraints were applied within the chain. Figure 8a) depicts a representative snapshot of the conformation obtained from such “control simulations”. We see that all the macrodomains are now well mixed. They do not possess any localisation and the polymer conformation is purely entropy-driven(6).

**Figure 8.**
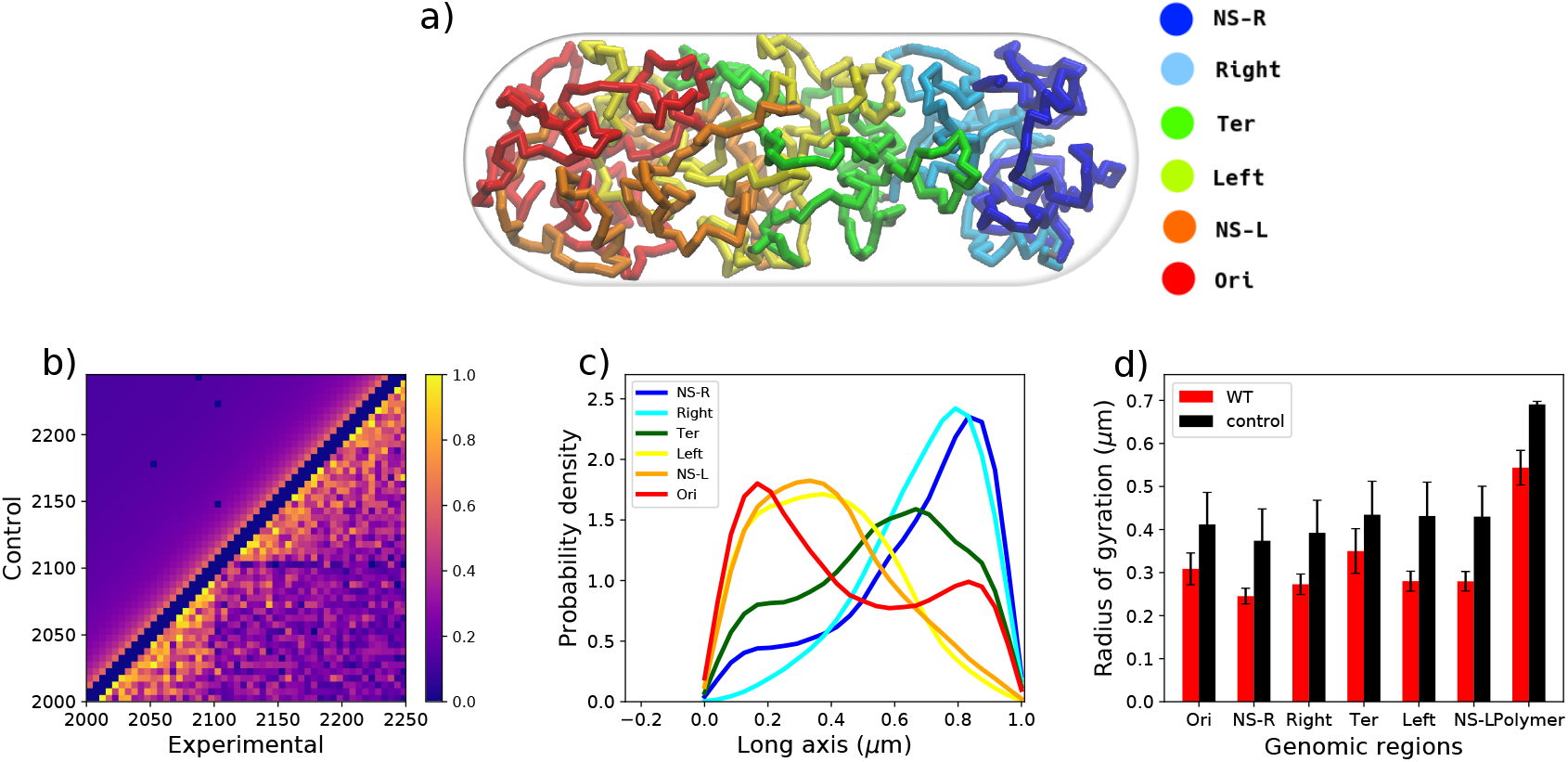
**a)** A color coded snapshot of the random-walk chromosome(control). **b)** Comparison of experimental and the control’s contact probability matrices. The matrices have been zoomed into a small section of the diagonal for clarity in comparison. **c)** Average densities of the four macrodomains and two Non-structured regions with respect to the cell long axis. **d)** Comparison of macrodomain sizes between control and WT.

We then generated the simulated Hi-C matrix for the control simulations. We see that the signature patterns along the diagonal, as observed in the original model (i.e. one modelled with Hi-C data) and experiment, are completely absent in Hi-C matrix obtained in this ‘control simulation’(Figure 8b). These patterns correspond to CIDs on a smaller scale, and macrodomains on a larger genomic scale. This shows that the contacts which are important, are relatively high in probability and likely arise due to presence of nucleoid-associated protein. The potential loss of the information in a chromosome model devoid of Hi-C contact probabilities data (as in the “control simulation”) can be further gleaned from Figure 8c. We see that the relative positions of the macrodomains are not correct (eg. Ori is at the pole). Most macrodomains have become disorganised since the distribution is more spread out, than in WT, with significant domain overlaps(Table S2). We see that all macrodomains have the same size in model derived from ‘control simulation’, suggesting that the localized interactions are missing.

## CONCLUSION

In conclusion, we report a Hi-C data-integrated comprehensive model of *E. coli* chromosome at 5 kbp resolution. The model is able to recover and represent the extent of information Hi-C encodes. The model captures the macrodomain segregation of chromosome precisely. The approach presented in the current work is distinct from other reconstruction algorithms recently used for modelling chromosomes(42, 43, 44) as the structures obtained from these algorithms do not produce an ensemble of structures and also requires scaling by another experimental data, such as FISH, to have the conformations commensurate with cell-sizes. On the contrary, the current model quantitatively reconciles numerous, independent experimental measurements on *E. coli* such as distances measured from FISH(7), experimental Recombination assay percentages(9) and linear densities(32). The model predicts a roughly linear organization of chromosome regions, in line with experimental investigation(10). The model predicts that in oriC and diff are located at the mid-cell diametrically opposite to each other in exponentially dividing cells which was also seen from experiments(31).The model also demonstrates that oriC localises at the mid-cell in most of the conformations (Figure 4c and d), consistent with the predictions the plectonemic model proposed by *Hacker et. al* where oriC was located at the mid-cell. We also were able to predict CID boundaries and the location of rrn operons using an indigenous way of analysing of radius of gyration of the chromosome segments. All these results reflect upon the multitude of information Hi-C already encodes and our model being able to capture them properly. We claim that the protocol for conformation generation is simple and fast with a high efficiency in reproducing experimental observations.

Taken together, the model brings out the multiscale and multi-faceted organization of bacterial chromosome, manifesting a helical, macrodomain-segregated morphology at large scale and CIDs at a fine-grained scale. Finally, a control model, which does not incorporate Hi-C data, shows that the multiscale organization and domain-segregation do not appear in such model. NAPs regulate these two factors for proper growth of the bacteria. The role of NAPs in maintaining overall chromosome conformation remains to be explored. Though we did not investigate the dynamics of the chromosome, but by incorporating mass of each bead one can, in practice, explore the chromosomal dynamics. Currently our model does not include replication arms, but its incorporation would yield a more precise picture of the *E. coli* chromosome. Incorporation of other experimental data would act as refinements on the ensemble averaged Hi-C which has been used as the basis for the modelling. multiscale simulations can also be attempted in which the coarse-grained interactions can be designed the way we incorporated bonds using Hi-C.

## ACKNOWLEDGEMENTS

This work was supported by computing resources obtained from shared facility of TIFR Centre for Interdisciplinary Sciences, India. JM would like to acknowledge research intramural research grants obtained from TIFR, DAE, India, Ramanujan Fellowship and Core Research grants provided by the Department of Science and Technology (DST) of India (CRG/2019/001219).

## REFERENCES

1. G Reshes, S Vanounou, I Fishov, and M Feingold. Timing the start of division in E. coli: a single-cell study. Physical biology, 5(4):046001, 2008.

2. Benjamin Volkmer and Matthias Heinemann. Condition-dependent cell volume and concentration of Escherichia coli to facilitate data conversion for systems biology modeling. PloS one, 6(7), 2011.

3. Mathew Stracy, Christian Lesterlin, Federico Garza De Leon, Stephan Uphoff, Pawel Zawadzki, and Achillefs N Kapanidis. Live-cell superresolution microscopy reveals the organization of RNA polymerase in the bacterial nucleoid. Proceedings of the National Academy of Sciences, 112(32):E4390–E4399, 2015.

4. Esteban Toro and Lucy Shapiro. Bacterial chromosome organization and segregation. Cold Spring Harbor perspectives in biology, 2(2):a000349, 2010.

5. Thøger J Krogh, Jakob Møller-Jensen, and Christoph Kaleta. Impact of chromosomal architecture on the function and evolution of bacterial genomes. Frontiers in microbiology, 9:2019, 2018.

6. Jagannath Mondal, Benjamin P Bratton, Yijie Li, Arun Yethiraj, and James C Weisshaar. Entropy-based mechanism of ribosome-nucleoid segregation in E. coli cells. Biophysical journal, 100(11):2605–2613, 2011.

7. Olivier Espéli, Romain Mercier, and Frédéric Boccard. DNA dynamics vary according to macrodomain topography in the E. coli chromosome. Molecular microbiology, 68(6):1418–1427, 2008.

8. Hironori Niki, Yoshiharu Yamaichi, and Sota Hiraga. Dynamic organization of chromosomal DNA in Escherichia coli. Genes & Development, 14(2):212–223, 2000.

9. Michèle Valens, Stéphanie Penaud,Michèle Rossignol, François Cornet, and Frédéric Boccard. Macrodomain organization of the Escherichia coli chromosome. The EMBO journal, 23(21):4330–4341, 2004.

10. Paul A Wiggins, Keith C Cheveralls, Joshua S Martin, Robert Lintner, and Jané Kondev. Strong intranucleoid interactions organize the Escherichia coli chromosome into a nucleoid filament. Proceedings of the National Academy of Sciences, 107(11):4991–4995, 2010.

11. Nastaran Hadizadeh Yazdi, Calin C Guet, Reid C Johnson, and John F Marko. Variation of the folding and dynamics of the e scherichia coli chromosome with growth conditions. Molecular microbiology, 86(6):1318–1333, 2012.

12. Remus T Dame, Fatema-Zahra M Rashid, and David C Grainger. Chromosome organization in bacteria: mechanistic insights into genome structure and function. Nature Reviews Genetics, pages 1–16, 2019.

13. Mark A Umbarger, Esteban Toro, Matthew A Wright, Gregory J Porreca, Davide Bau, Sun-Hae Hong, Michael J Fero, Lihua J Zhu, Marc A Marti-Renom, Harley H McAdams, et al. The three-dimensional architecture of a bacterial genome and its alteration by genetic perturbation. Molecular cell, 44(2):252–264, 2011.

14. Miriam Fritsche, Songling Li, Dieter W Heermann, and Paul A Wiggins. A model for Escherichia coli chromosome packaging supports transcription factor-induced DNA domain formation. Nucleic acids research, 40(3):972–980, 2012.

15. Julien Dorier and Andrzej Stasiak. Modelling of crowded polymers elucidate effects of double-strand breaks in topological domains of bacterial chromosomes. Nucleic acids research, 41(14):6808–6815, 2013.

16. Debasish Chaudhuri and Bela M Mulder. Molecular dynamics simulation of a feather-boa model of a bacterial chromosome. Methods Mol. Biol., pages 403–415, 2018.

17. Charlène Planchenault, Marine C Pons, Caroline Schiavon, Patricia Siguier, Jérôme Rech, Catherine Guynet, Julie Dauverd-Girault, Jean Cury, Eduardo PC Rocha, Ivan Junier, et al. Intracellular positioning systems limit the entropic eviction of secondary replicons toward the nucleoid edges in bacterial cells. Journal of Molecular Biology, 432(3):745–761, 2020.

18. David S Goodsell, Ludovic Autin, and Arthur J Olson. Lattice models of bacterial nucleoids. The Journal of Physical Chemistry B, 122(21):5441–5447, 2018.

19. Jing Huang and Tamar Schlick. Macroscopic modeling and simulations of supercoiled DNA with bound proteins. The Journal of chemical physics, 117(18):8573–8586, 2002.

20. Chris A Brackley, Stephen Taylor, Argyris Papantonis, Peter R Cook, and Davide Marenduzzo. Nonspecific bridging-induced attraction drives clustering of DNA-binding proteins and genome organization. Proceedings of the National Academy of Sciences, 110(38):E3605–E3611, 2013.

21. William C Hacker, Shuxiang Li, and Adrian H Elcock. Features of genomic organization in a nucleotide-resolution molecular model of the Escherichia coli chromosome. Nucleic acids research, 45(13):7541–7554, 2017.

22. Virginia S Lioy, Axel Cournac, Martial Marbouty, Stéphane Duigou, Julien Mozziconacci, Olivier Espéli, Frédéric Boccard, and Romain Koszul. Multiscale structuring of the E. coli chromosome by nucleoid-associated and condensin proteins. Cell, 172(4):771–783, 2018.

23. Tung BK Le, Maxim V Imakaev, Leonid A Mirny, and Michael T Laub. High-resolution mapping of the spatial organization of a bacterial chromosome. Science, 342(6159):731–734, 2013.

24. Marie Trussart, Eva Yus, Sira Martinez, Davide Bau, Yuhei O Tahara, Thomas Pengo, Michael Widjaja, Simon Kretschmer, Jim Swoger, Steven Djordjevic, et al. Defined chromosome structure in the genome-reduced bacterium Mycoplasma pneumoniae. Nature communications, 8(1):1–13, 2017.

25. Erez Lieberman-Aiden, Nynke L Van Berkum, Louise Williams, Maxim Imakaev, Tobias Ragoczy, Agnes Telling, Ido Amit, Bryan R Lajoie, Peter J Sabo, Michael O Dorschner, et al. Comprehensive mapping of long-range interactions reveals folding principles of the human genome. Science, 326(5950):289–293, 2009.

26. Asli Yildirim and Michael Feig. High-resolution 3D models of caulobacter crescentus chromosome reveal genome structural variability and organization. Nucleic acids research, 46(8):3937–3952, 2018.

27. Axel Cournac, Hervé Marie-Nelly, Martial Marbouty, Romain Koszul, and Julien Mozziconacci. Normalization of a chromosomal contact map. BMC genomics, 13(1):436, 2012.

28. Michele Di Pierro, Bin Zhang, Erez Lieberman Aiden, Peter G Wolynes, and José N Onuchic. Transferable model for chromosome architecture. Proceedings of the National Academy of Sciences, 113(43):12168–12173, 2016.

29. Saeed Saberi and Eldon Emberly. Chromosome driven spatial patterning of proteins in bacteria. PLoS computational biology, 6(11), 2010.

30. Mark James Abraham, Teemu Murtola, Roland Schulz, Szilard Pall, Jeremy C. Smith, Berk Hess, and Erik Lindahl. Gromacs: High performance molecular simulations through multi-level parallelism from laptops to supercomputers. SoftwareX, 1-2:19–25, 2015.

31. Xindan Wang, Xun Liu, Christophe Possoz, and David J Sherratt. The two Escherichia coli chromosome arms locate to separate cell halves. Genes & development, 20(13):1727–1731, 2006.

32. Somenath Bakshi, Albert Siryaporn, Mark Goulian, and James C Weisshaar. Superresolution imaging of ribosomes and RNA polymerase in live Escherichia coli cells. Molecular microbiology, 85(1):21–38, 2012.

33. Romain Mercier, Marie-Agnès Petit, Sophie Schbath, Stephane Robin, Meriem El Karoui, Frédéric Boccard, and Olivier Espéli. The matp/mats site-specific system organizes the terminus region of the E. coli chromosome into a macrodomain. Cell, 135(3):475–485, 2008.

34. Konstantin Klenin and Jörg Langowski. Computation of writhe in modeling of supercoiled DNA. Biopolymers: Original Research on Biomolecules, 54(5):307–317, 2000.

35. A Yu Grosberg, Sergei K Nechaev, and Eugene I Shakhnovich. The role of topological constraints in the kinetics of collapse of macromolecules. Journal de physique, 49(12):2095–2100, 1988.

36. Leonid A Mirny. The fractal globule as a model of chromatin architecture in the cell. Chromosome research, 19(1):37–51, 2011.

37. Alexander J Taylor and other SPOCK contributors. pyknotid knot identification toolkit. https://github.com/SPOCKnots/pyknotid, 2017. Accessed 2020-03.

38. Pauli Virtanen, Ralf Gommers, Travis E. Oliphant, Matt Haberland, Tyler Reddy, David Cournapeau, Evgeni Burovski, Pearu Peterson, Warren Weckesser, Jonathan Bright, Stéfan J. van der Walt, Matthew Brett, Joshua Wilson, K. Jarrod Millman, Nikolay Mayorov, Andrew R. J. Nelson, Eric Jones, Robert Kern, Eric Larson, CJ Carey, lİhan Polat, Yu Feng, Eric W. Moore, Jake Vand erPlas, Denis Laxalde, Josef Perktold, Robert Cimrman, Ian Henriksen, E. A. Quintero, Charles R Harris, Anne M. Archibald, Antônio H. Ribeiro, Fabian Pedregosa, Paul van Mulbregt, and SciPy 1. 0 Contributors. SciPy 1.0: Fundamental Algorithms for Scientific Computing in Python. Nature Methods, 17:261–272, 2020.

39. Julio E Cabrera and Ding J Jin. Active transcription of rRNA operons is a driving force for the distribution of RNA polymerase in bacteria: effect of extrachromosomal copies of rrnb on the in vivo localization of RNA polymerase. Journal of bacteriology, 188(11):4007–4014, 2006.

40. Scott A Scholz, Rucheng Diao, Michael B Wolfe, Elayne M Fivenson, Xiaoxia Nina Lin, and Peter L Freddolino. High-resolution mapping of the Escherichia coli chromosome reveals positions of high and low transcription. Cell systems, 8(3):212–225, 2019.

41. Sandra Goetze, Julio Mateos-Langerak, Hinco J Gierman, Wim de Leeuw, Osdilly Giromus, Mireille HG Indemans, Jan Koster, Vladan Ondrej, Rogier Versteeg, and Roel van Driel. The three-dimensional structure of human interphase chromosomes is related to the transcriptome map. Molecular and cellular biology, 27(12):4475–4487, 2007.

42. Annick Lesne, Julien Riposo, Paul Roger, Axel Cournac, and Julien Mozziconacci. 3D genome reconstruction from chromosomal contacts. Nature methods, 11(11):1141, 2014.

43. Guillaume Le Treut, François Képès, and Henri Orland. A polymer model for the quantitative reconstruction of chromosome architecture from hic and gam data. Biophysical journal, 115(12):2286–2294, 2018.

44. Ahmed Abbas, Xuan He, Jing Niu, Bin Zhou, Guangxiang Zhu, Tszshan Ma, Jiangpeikun Song, Juntao Gao, Michael Q Zhang, and Jianyang Zeng. Integrating hi-c and fish data for modeling of the 3D organization of chromosomes. Nature communications, 10(1):1–14, 2019.

