## Supplementary material for "Mapping the Multiscale Organisation of Escherichia Coli Chromosome in a Hi-C-integrated Model": SI

### Supplementary Information

#### Calculation of scaling parameter and cell size

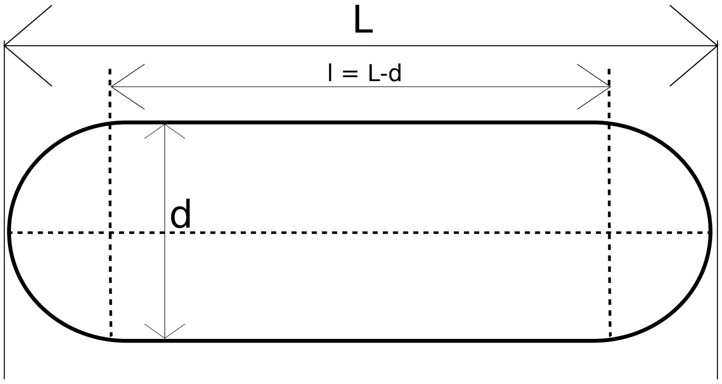


**Figure S1:** Cell dimensions

At 37°C in LB media, an estimated cell length (including end-caps) would be 2.482 $\mu$m while the diameter would be 0.933 $\mu$m [1].
Let the order of the HiC contact frequency matrix be N, length of the cell (including end caps) be L, diameter be d and size of each polymer bead be $\sigma$.
Therefore, the number of beads in the polymer = N
Let l = L - d, r = $\frac{d}{2}$
Therefore, volume of the cell = $\frac{4}{3}\pi r^3 + \pi r^2 l = \pi r^2(\frac{4}{3}r + l)$
Assuming a volume fraction $f_r$,

$$\text{the volume occupied by the DNA} = f_r \text{ x Volume of the cell} = \pi r^2 f_r(\frac{4}{3}r + l) \tag{S1}$$

$$\text{But the volume occupied by the DNA} = \frac{4}{3}\pi(\frac{\sigma}{2})^3 = \frac{\pi\sigma^3}{6} \tag{S2}$$

On comparing equations (S1) and (S2), we get

$$\sigma = \sqrt[3]{\frac{6f_r r^2}{N}(\frac{4}{3}r + l)} \tag{S3}$$

Using $f_r = 0.1$[2] and the cell dimensions stated above for a resolution of 5 kbp (N=928), $\sigma = 0.067348$ $\mu$m.
Upon setting $\sigma = 1$ and scaling the cell dimensions provided above, we get the cell length (including end-caps) equals to 36.884 and the diameter equals to 13.864 in reduced (simulation) units.

#### Generation of initial structures

We have used 200 randomly generated beads-on-a-spring polymer conformations for our simulations. Each conformation has 928 beads, which is equal to the number of bins the DNA has been divided into to have a resolution of 5 kbp, or also equal to the order of the HiC contact frequency matrix at 5 kbp. For a matrix with different order, hence resolution, one needs to generate polymer conformation(s) with the number of beads equal to its order. Each polymer conformation was grown bead by bead while maintaining no overlap between beads. Since the simulations would be performed inside a confinement mimicking the cell boundary, the beads were generated in such a way that the polymer beads stay within the cell boundaries even during the growth of the polymer. This was implemented so as to not cause huge force unbalances during energy minimization, thereby making the process faster.

#### Interaction potenitals

##### Finding a good transfer function

The main purpose of the current work is to develop a quantitative model that can explore the higher order organisation of the 4.64 Mbp long, *E. coli* chromosome. Towards the end, following the protocol outlined in Figure 1 and explained in *Methods*, we integrate recently reported Hi-C interaction matrix of *E. coli* chromosome [3] within a polymer based beads-on-a-spring model. The chromosome, thereby modelled, is subsequently subjected to a langevin dynamics simulation at a friction coefficient of 1 $\sqrt{\frac{m\epsilon}{\sigma}}$ and temperature of 303K. The simulation observables are statistically averaged over the last 2000 frames from the 200 trajectories.

In previous attempts to model the chromosome using Hi-C data[3, 4], a variant of the inverse function has been used to convert contact probabilities to distances(S4).

$$D \propto P^{-s} \tag{S4}$$

where $D$ is the distance matrix and $P$ is the contact probability matrix. The operation is elementwise inverse of matrix elements of $P$ raised to a power $s$. $s$ can be obtained by comparing the distances obtained by varying $s$ to an experimentally obtained set of distances[3]. We have used $s = 1.0$ for our studies[4] with a proportionality constant of $\sigma$ which is the bead size. Thus when the contact probability is 1, the distance of a Hi-C bond is $\sigma$. An earlier approach to model the bacterial chromosome using Hi-C data involved a low, but constant, force constants for all the contacts that are put into the model[5]. Here we scaled the force constants with respect to the distances, which in turn makes the force constants scale with contact probabilities. We hypothesize that if a pair of chromosome regions have a high contact proability between them, then they have been actively brought together and appear as a 'contact' in most cells when Hi-C is performed. Whereas regions with low contact probabilities are a result of the random collisions between different regions of the chromosome. To mimic this stochastic behaviour, we used a function, which we call the "transfer function", that would scale in such a way so that the lower the contact probability, lower is the force constant between two such regions. Thus for regions with no contact, there would not be any restraint or a restraint with a realtively very weak force constant, thereby introducing random fluctuations in their equlibrium distances.

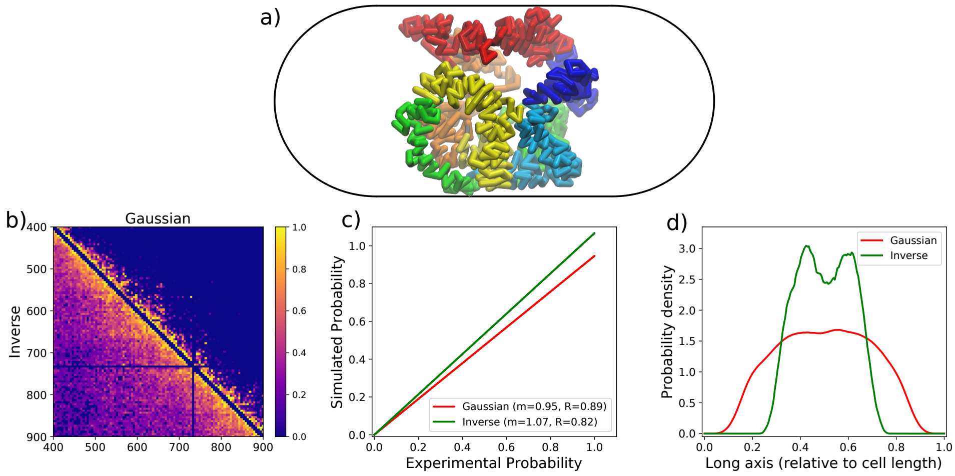


**Figure S2: a)** A snapshot of the 3D conformation for simulations using Eq. (S5). **b)** Comparison between the force constant matrices for Eqs. (**1**) and (S5). The lower half of the plot is for Eq. (S5) while the upper half is for Eq. (**1**). **c)** Compairson of the Hi-C simulated matrices using Eqs. (**1**) and (S5). **d)** Linear density along the long axis of the cell for simulations using Eqs. (**1**) and (S5).

First we tried a simple inverse transfer function as shown in Eq. (S5). $k_0$ is a parameter and does not effect the chromosome organization in anyway. It controls the maximum strength of the "Hi-C" bonds between two beads, thus can be chosen to arbitrarily low with respect to the adjacent beads' bond strength. For all simulations, $k_0 = 10$ was used.

$$k_{ij} = \frac{k_0}{D_{ij}} \tag{S5}$$

Though the distances we obtained correlated well with existing experimental data (Figure S4), but the chromosome was found to be very condensed inside the cell (Figure S2a), which is not in agreement with the linear density profiles of the chromosome from experiments and theoretical studies[6, 7]. Not to mention that it is difficult to deal with very low probabilities which get converted to very large distances. We had to define a distance cutoff (diameter of the cell), beyond which we did not consider them as contacts anymore. But the choice was arbitrary.

After using a simple inverse function, we tried a gaussian function to get the force constants (Eq. (S7)). The gaussian ensured that the at infinites and large distances, the force constant scaled down to such a low value that it can be safely ignored. Also, only a small fraction of the total number of contacts were actually used when we used Eq. (S7) for simulations as can be seen from Figure S2b and Figure S3. We see that when we use the gaussian, the important regions of contacts are mainly located along the diagonal and the small patch on the ends of the off-diagonal due to circularity of the chromosome. We see that such a small number of contacts is able to predict the whole contact parobbaility matrix with a high accuracy (Figure S2c). We also see that predictions are better when we used Eq. (S7). The gaussian also lets the chromosme spread out into the cell volume much more than the inverse transfer function (Figure S2d). This is a more realistic image as seen from previous theoretical and experimantal studies[6, 7]. Thus we believe that Eq. (S7) is a much better transfer function to generate more plausible 3D conformations.

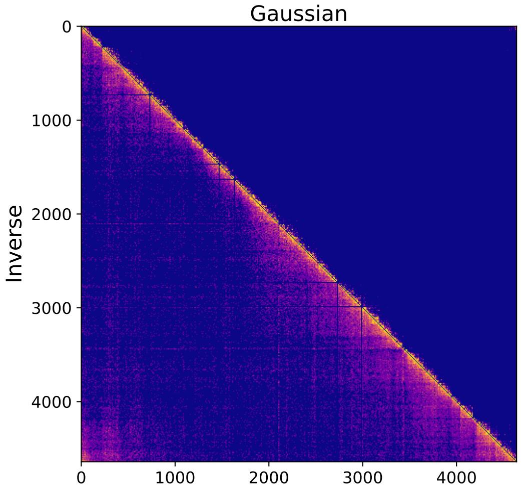


**Figure S3:** Comparison between the force constant matrices for Eqs. (**1**) and (S5) in the main text. The lower half of the plot is for Eq. (S5) while the upper half is for Eq. (**1**) in the main text.

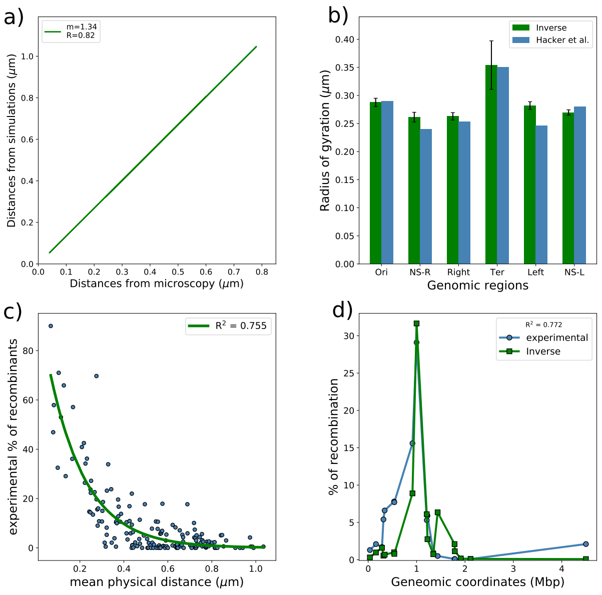


**Figure S4:** **a)** Comparison of distances measured via FISH[8] with distances obtained with simulations using inverse transfer function. **b)** Comparison of radii of gyration among domains with respect to Hacker *et. al*'s reported values for oriC@midcell in their plectonemic model[9]. **c)** Comparison between the recombination frequencies provided by *Valens et al. 2004* [10] vs. mean physical distance between recombination loci and the green line indicates the single exponential fit. **d)** A representative plot of recombination frequencies predicted by mean physical distance.
**blue: experimental data. green: predicted by simulated data using inverse transfer function.**

#### Bonded interactions

Adjacent beads of the polymer are connected by strong (300 kJ mol$^{-1}\sigma^2$) harmonic springs with $\sigma$ as the equilibrium bond length. This has been implemented by introducing harmonic force fields between adjacent beads. HiC contacts have also been modelled

as harmonic springs but with variable strengths and variable bond lengths.
Let the HiC contact probability matrix be $P$ and a distance matrix, $D$, has been defined such that

$$D_{ij} = \frac{\sigma}{P_{ij}} \tag{S6}$$

where $ij$ suggests the element in the i$^{th}$ row and j$^{th}$ column of the matrices. It should be noted that the matrix $P$ is a sparse matrix. Therefore a lot of the elements in D would be $\infty$. They would be taken care by the model itself as discussed below. The force constants for the bonds incorporating the HiC contacts into the model is given by

$$k_{ij} = k_0 e^{-\frac{(D_{ij}-\sigma)^2}{w}} \tag{S7}$$

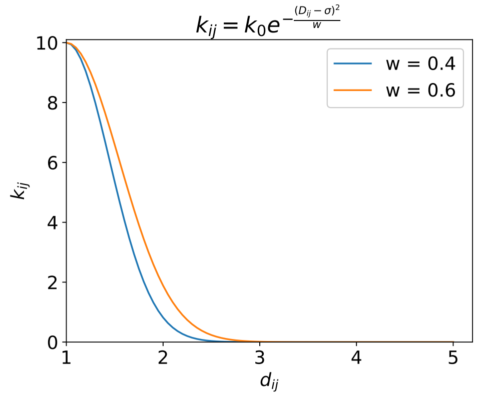


**Figure S5:** $k_{ij}$ vs. $D_{ij}$ (eqn (5)). $\sigma$=1.

As per Hooke's law,

$$\begin{aligned} V_{HiC}(D_{ij}) &= \frac{1}{2}k_{ij}(D_{ij} - r_{ij})^2 \\ \implies \quad V_{HiC}(D_{ij}) &= \frac{1}{2}k_0 e^{-\frac{(D_{ij}-\sigma)^2}{w}}(D_{ij} - r_{ij})^2 \end{aligned}$$

For $D_{ij} = \infty$, $V_{HiC}(D_{ij}) = 0$.

In equation (5), $k_0$ and $w$ are parameters that need to be optimised but $k_0$ is just a amplitude term that determines the upper limit to the force constants of the "HiC bonds". Since a high force constant makes bonds rigid, a very high value of $k_0$ will only effect the net dynamics of the polymer. $k_0$ does not in any form effect the equilibrium distances of the HiC bonds, thus has negligible effect on the overall polymer configuration. For all our simulations we used $k_0 = 10$. This choice is somewhat arbitrary, but the value of $k_0$ is low when compared to the force constant of the backbone bonds. This is not arbitrary. The reason for such a choice is that we have used contact **probability** to model these bonds. Therefore, a non-zero number of times it should be seen that there exists no HiC-contact between beads $i$ and$j$ if we sample from a large set of time series

distances at equilibrium between $i$ and $j$. A low force constant would lead to larger deviations in the distance between the two beads during simulations and should mimic the sampling, i.e. if we look at the probability distribution of the time series distance data of the HiC bond length between two beads $i$ and $j$, we shall see that the equilibrium would have a probability almost equal to $P_{ij}$.
$w$ needs to be optimized for each unique HiC contact probability matrix. It determines the width of the Gaussian in equation (5), therefore behaves as a filtering parameter. For example, from Fig-2 it can be seen that for a higher value of $w$, more number of $k_{ij}$ have magnitudes greater than 0. This incorporates more HiC bonds into the system. To speed up simulations, we did not consider bonds whose force constants were lower than $10^{-7}$. Such bonds are very weak and do not contribute significantly to the dyamics of the chromosome.

#### The cell boundary

The cell boundary has been implemented by using a restraining potential of the form

$$V_{res}(r; R_0) = \frac{1}{2} k_{res} |\vec{r} - \vec{R_0}|^2 \ \mathbf{H}(|\vec{r} - \vec{R_0}|) \tag{S8}$$

$\mathbf{H}$ is a step function and activates if a particle goes out of the confinement, which, here is a spherocylinder. $R_0$ is the center of the spherocylinder. $k_{res}$ determines how rigid is the cell wall. For simulations we have used 310 kJ mol$^{-1}\sigma^2$, slightly higher than $k_{adj}$, making it the highest in magnitude among all the force constants present.

#### Simulation details

For each cell type we have discussed, 200 simulations have been performed with different initial structures. Each simulation consisted of 2 steps: i) An energy minimization. ii) A production run using stochastic dynamic simulations.
For both of the above steps, HiC interactions were considered. The time step for step (ii) was 2 fs. Each production run has been run for $2\times10^6$ steps from which only the last 2000 frames have been used for further analysis.

#### Generation of simulated contact probability matrix

Using the last 2000 frames of $k$-th simulation, we generate a distance matrix $D^k$ whose distances have been averaged over the number of frames. We generate the final simulated distance matrix $D_{\text{sim}}$ by averaging over those matrices.

$$D_{\text{sim}} = \frac{1}{n} \sum_{k=1}^{n} D^k \tag{S9}$$

Similarly, we also generate $P^k$ which is the simulated contact probability matrix for the $k$-th simulation.

$$P^k_{ij} = \frac{\sigma}{D^k_{ij}} \tag{S10}$$

The simulated probability matrix $P_{\text{sim}}$ is an average over $P^k$.

$$P_{\text{sim}} = \frac{1}{n} \sum_{k=1}^{n} P^k \tag{S11}$$

We have used n=200, i.e. we have used 200 simulations for the averaging.
It should be noted that $P_{\text{sim}} \neq \frac{\sigma}{D_{\text{sim}}}$, and it is evident from the definitions of $P^k$, $D^k$, $P_{\text{sim}}$, $D_{\text{sim}}$ given by equations (7) - (9).
Each frame corresponds to a microstate of the ensemble of chromatin conformations whose average is the experimental HiC matrix. Since a set of 2000 frames belong to a particular initial conformation, instead of generating 200 x 2000 = 4 x $10^5$ matrices and performing an average over them, we average over 200 matrices, which have been averaged frame-wise. We also generate a distance matrix averaged over frames and simulations. This matrix is used for all distance related calculations or analysis.

#### Filtering of simulated contact probability matrices

A lot of rows and columns in the experimental contact probability matrix are zero. This can be due to two reasons: i) there is no contact between those two regions. ii) the region is too compact for Hi-C to detect any contact. Since they are zero, after incorporating into the model, "collisions" between such regions are purely random, even they might be highly dense. This can lead to wrong interpretation of the data. To avoid this, we filter the simulated contact probability matrix by replacing elements in the matrix with zeros which are also zero in the experimental matrix. Thus we cannot get more information than what the experimental holds, but misinterpretations can be avoided. Solely for the purpose of having a better contrast within a matrix, we have raised each element to a power of 0.3 , only during plotting, so as to enhance the smaller values. For all numerical analysis we have used non-enhanced matrices.

#### Comparison between matrices

To compare matrices, we first *flatten* the matrices. Flattening simply means we convert the matrix into an 1D array, where each consecutive row adds to the array after the previous row. This array would thus be n x n long for a contact probability matrix of order n. Since the values which are zero in the experimental matrix are already zero, we recreate an array without any zeros. Two such arrays, without any zeros are compared for correlation or linear fitting. For absolute difference analysis also, we do not consider the zeros.

#### Optimization of $w$

The metric used to optimize $w$ is a Pearson correlation between the experimental and the *filtered* simulated contact probability matrices. Many elements in the contact probability matrix are zero due experimental limitations and experimental protocol. But in the simulated HiC matrix, we do not have any zero, but a base value of contact probability(since no two bead can go beyond L). Thus to make them comparable we set those values to zero the simulated matrix which are also zero in the experimental matrix. This is what we refer to when we mention filtered/filtering. Then the matrices are "flattened" in the sense that the 2D sparse matrix is converted to a 1D array with no non-zero elements. The zeros are removed so as to get the Pearson correlation between the non-zero elements only. Considering the zeros after filtering would have contributed to a higher value of the correlation coefficient, thereby reducing its sensitivity to $w$. In this way we are able to perform an element wise comparison between the matrices.

#### Non-bonded interactions

Since no information is known about the attractive interactions between DNA beads at 5 kbp resolution, we assume that all the attractive interactions bringing regions of the DNA close to each other have been captured by HiC. Therefore the non-bonded interactions are purely repulsive and are given by

$$V_{nb}(r) = \frac{A}{r^{12}} \tag{S12}$$

where A $= 4\epsilon\sigma^{12}$. Equation (6) is simply the repulsive part of the Lennard-Jones potential. This also ensures that polymer beads do not overlap significantly when equilibration is performed. For the simulations, A=1.0 has been used.

#### Simulation engine

We have used a modified GROMACS 5.0.7 as our simulation engine. The reduced units are as per the softwares specifications. Mass of each bead was set to 1 amu (atomic mass unit). The length scale unit used for simulations is nm ($\sigma = 1$ nm). Thus the cell dimensions become 36.884 nm and 13.86 nm. The temperature of the system was set to 303K and the friction was 2 amu/ps. Since the effect of temperature on distances should already have been captured by Hi-C, one can use any temperature around 310K if one does not need to explore the dynamics. The initial configurations were prepared as per GROMOS96 file format or a .gro file.

In the version of the software used, parallel simulations did not work without pbc (there should be no pbc since we are simulating one single molecule). To tackle this, we put our system at the center of a box ten times (arbitrarily chosen) bigger than the system boundaries. Then we performed productions runs with pbc implemented. Since the polymer cannot get out of the confinement, it will never cross the pbc. This allowed us to perform the production runs in parallel thereby producing a major speed up in simulation time.

#### Rendering 3D conformations

We have used the open-source package Visual Molecular Dynamics (VMD)[11] to render the representative 3D conformations of the chromosome model.

#### Crossing number and Writhe

The terms crossing number and writhe for polymers, such as the chromosome, comes from Knot Theory. In knot theory, the definition of the average crossing number is the number of times a knot diagram (3D coordinates of a closed curved with or without knots) crosses itself when projected onto a plane, averaged over all possible projections. Each crossing is assigned a sign depending upon whether it crosses over (+) or under (-) (Figure S6 a and b). The sum of the signed crossings provides the writhe (Figure S6c). It is a measure of how coiled a conformation is.

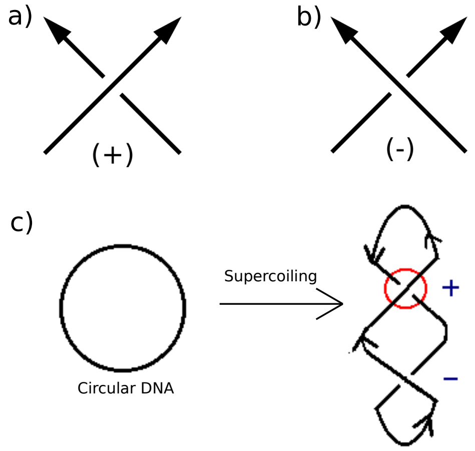


**Figure S6: a)** An over crossing[12]. **b)** An under crossing[12]. **c)** The circular DNA forms a helix upon supercoiling but with equal probabilities of positive and negative crossing, thus a net zero writhe[13].

### Supplementary Tables

**Table S1:** Average radius of gyration ($R_g$) and end-to-end distances ($R_e$) of all macrodomains, non-structured regions and the whole chromosome in $\mu$m.

| Variable | NS-R | Right | Ter | Left | NS-L | Ori | Chromosome |
|---|---|---|---|---|---|---|---|
| $R_g$ | 0.245±0.018 | 0.273±0.024 | 0.350±0.052 | 0.280±0.023 | 0.279±0.023 | 0.308±0.037 | 0.543±0.040 |
| $R_e$ | 0.501±0.006 | 0.540±0.006 | 0.904±0.007 | 0.661±0.006 | 0.623±0.006 | 0.676±0.006 | 1.573±0.448 |

**Table S2:** Macrodomain overlap scores

| MD1 | MD2 | WT | control |
|---|---|---|---|
| NS-R | Right | 0.416±0.123 | 0.656±0.188 |
| NS-R | Ter | 0.323±0.129 | 0.464±0.236 |
| NS-R | Left | 0.210±0.166 | 0.368±0.248 |
| NS-R | NS-L | 0.168±0.149 | 0.323±0.234 |
| NS-R | Ori | 0.316±0.108 | 0.348±0.270 |
| Right | Ter | 0.461±0.100 | 0.592±0.164 |
| Right | Left | 0.165±0.155 | 0.402±0.223 |
| Right | NS-L | 0.113±0.138 | 0.336±0.232 |
| Right | Ori | 0.153±0.155 | 0.327±0.248 |
| Ter | Left | 0.372±0.124 | 0.642±0.174 |
| Ter | NS-L | 0.182±0.184 | 0.460±0.226 |
| Ter | Ori | 0.221±0.177 | 0.386±0.257 |
| Left | NS-L | 0.419±0.117 | 0.619±0.166 |
| Left | Ori | 0.251±0.177 | 0.478±0.252 |
| NS-L | Ori | 0.389±0.122 | 0.720±0.183 |

### Supplementary Figures

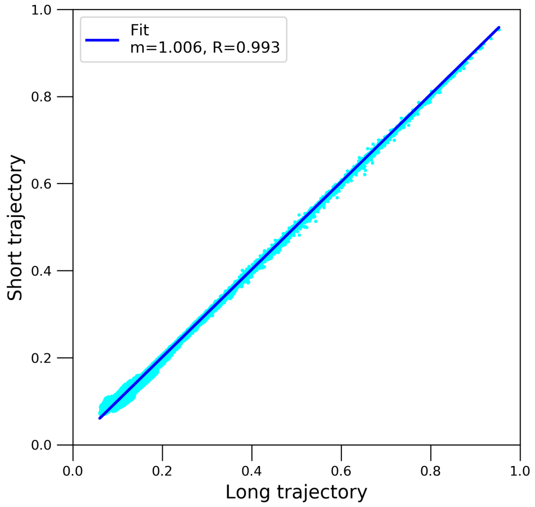


**Figure S7:** Comparison between simulated Hi-C matrices for short($2x10^6$) and long($12x10^6$) simulations for wildtype *E. coli* at 37°C. The deep blue line is the linear fit.

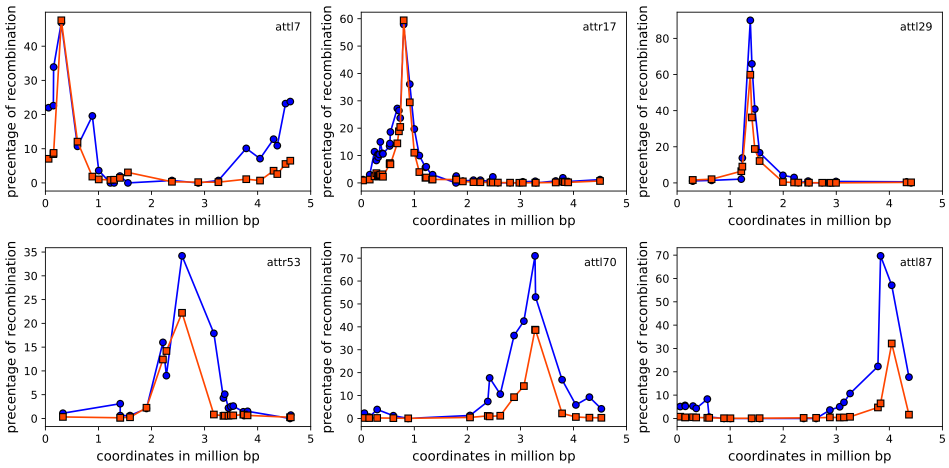


**Figure S8:** Predicted recombination frequency based on the exponential curve fitted in the figure 5 for 6 other loci (attl7, attr17, attl29, attr53, attl70, and attl87). Orange line indicates the fitted data and blue is experimentally observed data from *Valens et. al. (2004)*[10].

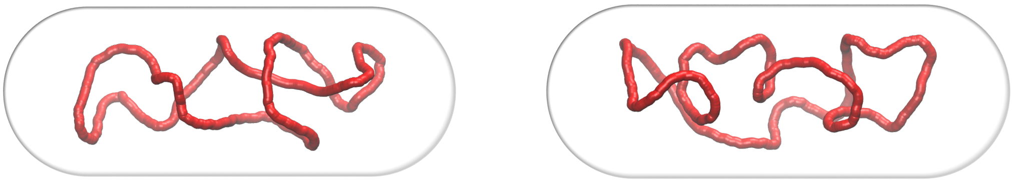

**Figure S9:** Both are representative snapshots of the COGs of the polymer calculated using a window size of 20.

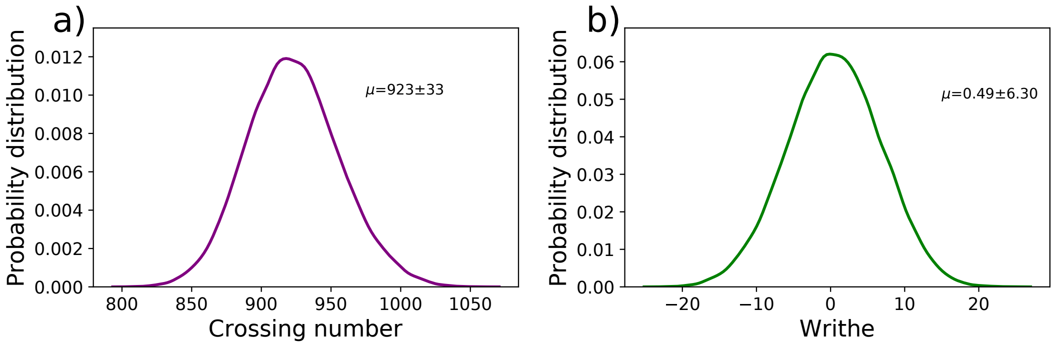


**Figure S10: a)**Distribution of number of crossings from different frames of the whole chromosome. **b)** Distribution of writhe from different frames of the whole chromosome.

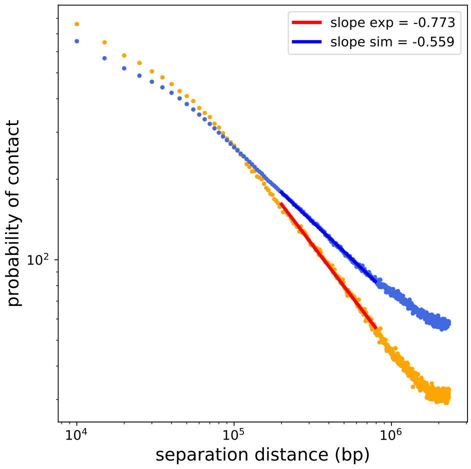


**(a)** Fractal globule plot

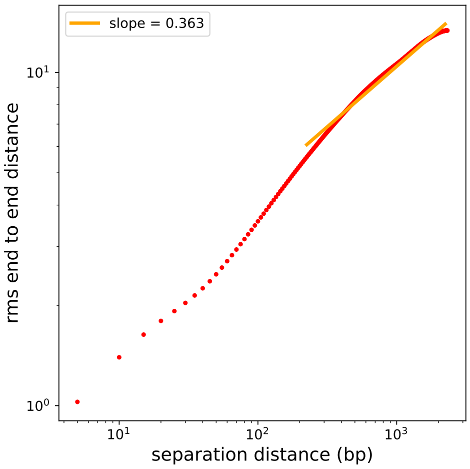


**(b)** Rms end-to-end distance plot

**Figure S11: a)** fractal globule plot for experimental and simulated contact probability matrix (orange: experimental, blue: simulated), **b)** mean of rms end-to-end distance of the genomic segments (in increasing length of 5000bp) vs. genomic distance (bp).

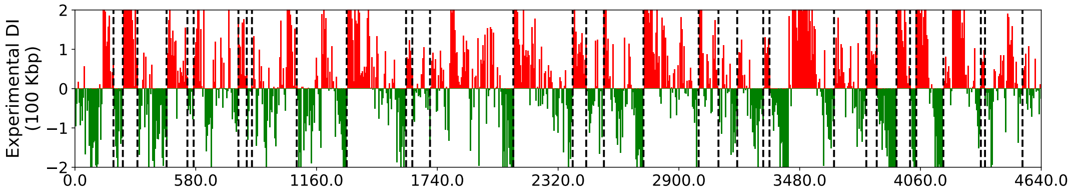


**Figure S12:** CID boundaries calculated using DI on the experimental contact probability matrix.

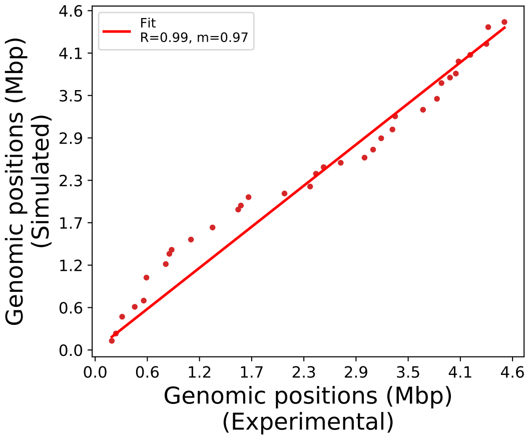


**Figure S13:** Correlation between CID boundaries detected by $R_g$ map and DI

### References


[1] G Reshes, S Vanounou, I Fishov, and M Feingold. Timing the start of division in E. coli: a single-cell study. *Physical biology*, 5(4):046001, 2008.

[2] Saeed Saberi and Eldon Emberly. Chromosome driven spatial patterning of proteins in bacteria. *PLoS computational biology*, 6(11), 2010.

[3] Virginia S Lioy, Axel Cournac, Martial Marbouty, Stéphane Duigou, Julien Mozziconacci, Olivier Espéli, Frédéric Boccard, and Romain Koszul. Multiscale structuring of the E. coli chromosome by nucleoid-associated and condensin proteins. *Cell*, 172(4):771–783, 2018.

[4] Annick Lesne, Julien Riposo, Paul Roger, Axel Cournac, and Julien Mozziconacci. 3D genome reconstruction from chromosomal contacts. *Nature methods*, 11(11):1141, 2014.

[5] Asli Yildirim and Michael Feig. High-resolution 3D models of caulobacter crescentus chromosome reveal genome structural variability and organization. *Nucleic acids research*, 46(8):3937–3952, 2018.

[6] Jagannath Mondal, Benjamin P Bratton, Yijie Li, Arun Yethiraj, and James C Weisshaar. Entropy-based mechanism of ribosome-nucleoid segregation in E. coli cells. *Biophysical journal*, 100(11):2605–2613, 2011.

[7] Somenath Bakshi, Albert Siryaporn, Mark Goulian, and James C Weisshaar. Super-resolution imaging of ribosomes and RNA polymerase in live Escherichia coli cells. *Molecular microbiology*, 85(1):21–38, 2012.

[8] Olivier Espéli, Romain Mercier, and Frédéric Boccard. DNA dynamics vary according to macrodomain topography in the E. coli chromosome. *Molecular microbiology*, 68(6):1418–1427, 2008.

[9] William C Hacker, Shuxiang Li, and Adrian H Elcock. Features of genomic organization in a nucleotide-resolution molecular model of the Escherichia coli chromosome. *Nucleic acids research*, 45(13):7541–7554, 2017.

[10] Michèle Valens, Stéphanie Penaud, Michèle Rossignol, François Cornet, and Frédéric Boccard. Macrodomain organization of the Escherichia coli chromosome. *The EMBO journal*, 23(21):4330–4341, 2004.

[11] William Humphrey, Andrew Dalke, and Klaus Schulten. VMD – Visual Molecular Dynamics. *Journal of Molecular Graphics*, 14:33–38, 1996.

[12] Wikipedia contributors. Writhe — Wikipedia, the free encyclopedia, 2020. [Online; accessed 4-September-2020].

[13] S. Harrell M. Beals, L. Gross. DNA and knot theory. http://www.tiem.utk.edu/~gross/bioed/webmodules/DNAE>coknot.html, 1999. Accessed 2020-08.
